# NAND Hybrid Riboswitch Design by Deep Batch Bayesian Optimization

**DOI:** 10.1101/2025.03.28.645907

**Authors:** Daniel Kelvin, Erik Kubaczka, Heinz Koeppl, Beatrix Suess

**Author notes:** Correspondence can be addressed to and. These authors contributed equally (Daniel Kelvin, Erik Kubaczka).

## Abstract

The design of large genetic circuits requires genetic regulatory devices capable of performing complex logic operations. Hybrid riboswitches, synthetically enhanced compact RNA elements (<100 nucleotides) that form a tertiary structure with the ability to specifically bind two different target molecules, can be used to design genetic regulators that emulate Boolean logic. When inserted into the 5’ UTR of an mRNA, these devices can regulate translation initiation upon specific binding of one or both ligands. The goal of this study is to design hybrid riboswitches that emulate Boolean NAND logic in yeast. We propose a novel machine learning-based design framework combining high-throughput *in vivo* screening and deep Bayesian optimization. Through an initial screening, we discovered a hybrid riboswitch with NAND behavior. Using batch Bayesian optimization with an ensemble neural network as surrogate, we further improve the NAND functionality of our hybrid riboswitch with respect to a performance score, thereby achieving near digital NAND behavior. With its focus on model-based and score-driven design, our proposed method can complement experiment driven approaches by allowing fine grained adaptation of functionality, including constructs sensitive to single nucleotide changes.

## Introduction

Information processing plays a key role in cellular pathways as well as synthetically engineered devices. To engineer information processing synthetically, constructs imitating Boolean logic in cells are ideal (Bressler et al., 2023; Chen et al., 2020; Mohammadi et al., 2017; Stanton et al., 2014). RNA based gene regulation, as implemented by riboswitches, provides an appealing basis for the design of logic devices. Riboswitches are RNA sequences with the ability to influence gene expression through the formation of complex tertiary structures that specifically bind target molecules (ligands) (Kavita and Breaker, 2023; Nahvi et al., 2002). As they require no auxiliary factors to regulate gene expression, riboswitches impose only a low metabolic burden on the host organism (Tucker and Breaker, 2005). While naturally occurring riboswitches have been discovered mostly in bacteria, synthetic riboswitches have been engineered to utilize novel regulatory mechanisms for implementation not only in bacteria but also in eukaryotes (Etzel and Mörl, 2017; Kavita and Breaker, 2023; McCown et al., 2017; Weigand and Suess, 2007; Wieland and Fussenegger, 2010). In *Saccharomyces cerevisiae,* synthetic riboswitches utilizing a roadblock mechanism prevent translation initiation by stabilizing their structure upon ligand binding (Hanson et al., 2003; Suess et al., 2003). To enable the design of constructs implementing multi-input Boolean logic, riboswitch architectures capable of accepting two or more inputs are required. Hybrid riboswitches, which contain two different ligand binding pockets in one continuous structure, meet this requirement (Kelvin et al., 2025). In addition to being excellent regulators, the ability to bind both ligands separately and simultaneously makes hybrid riboswitches suitable candidates for the implementation of Boolean logic. In particular, the hybrid riboswitch architecture has the potential to implement NAND behavior, a mechanistically challenging type of logic gate that has not been observed in any naturally occurring dual-input riboswitch (Sherlock et al., 2022). Previous attempts at engineering riboswitches to implement NAND behavior have not taken advantage of direct interactions between different binding pockets. Instead, they have relied on a ribozyme to act as a shared expression platform that is controlled by two separately inserted binding pockets (Win and Smolke, 2008; Felletti et al., 2016).

The development of a synthetic riboswitch that utilizes the roadblock mechanism begins with an *in vitro* selection process known as SELEX (Systematic Evolution of Ligands by EXponential Enrichment) (Tuerk and Gold, 1990; Ellington and Szostak, 1990; Nutiu and Li, 2005; Stoltenburg et al., 2012; Boussebayle et al., 2019). This method is used to discover sequences with high binding affinity for the chosen ligand, called aptamers. Subsequent *in vivo* screenings are required to identify candidates capable of undergoing conformational changes essential for mediating gene expression and to optimize their dynamic range. This approach focuses on binding affinity maximization and is not suitable for the design of devices with multiple inputs, such as hybrid riboswitches. Previously reported hybrid riboswitches have been created using rational design (Kelvin et al., 2025), which also poses a challenge when attempting to achieve specific switching behavior, due to the complex sequence-to-function relationship of the roadblock mechanism (Weigand et al., 2011). In addition, the sensitivity of (hybrid) riboswitches to single nucleotide changes makes the high-throughput generation of datasets containing regulatory active constructs for *in silico* approaches (Angenent-Mari et al., 2020; Valeri et al., 2020) inefficient. This lack of robustness poses a challenge in the design of multi-input riboswitches that require structural, sequence-dependent interactions between the different ligand binding pockets to emulate a Boolean switching pattern. Additionally, the method has to cope with the exponential complexity of the sequence space.

In modern machine learning, Bayesian optimization (Frazier, 2018; Garnett, 2023; Shahriari et al., 2016; Wang et al., 2022), a sequential optimization method suitable for costly to evaluate black box functions, is often used in the context of hyper parameter optimization for deep learning (Akiba et al., 2019; Hutter et al., 2019; Snoek et al., 2012), but has also been applied to biological engineering for the design of proteins (Wu et al., 2019; Yang et al., 2025, 2019), DNA (Friedman et al., 2025), small molecules (Bailey et al., 2024; Stanton et al., 2022), and experimental conditions (Dürholt et al., 2024; Fitzner et al., n.d.; Rosa et al., 2022). In a round based manner, new candidate RNA sequences for experimental evaluation are proposed and the gained information is taken into account in the next rounds. This compares to active learning (Settles, 2009; Gal et al., 2017; Di Fiore et al., 2024), with the difference in the criterion for candidate selection. Using a probabilistic surrogate model that accounts for uncertainty, classically a Gaussian process (Garnett, 2023; Quadrianto et al., 2010), Bayesian optimization can efficiently trade off exploration and exploitation, also in combinatorial design spaces such as RNA sequences. A promising trend -especially in life sciences- is the use of deep learning based surrogate models (Bailey et al., 2024; Friedman et al., 2025; Stanton et al., 2022; Yang et al., 2025, 2019), which are successfully applied as sequence-to-function models but require careful design for well calibrated uncertainty quantification. For the candidate selection, different acquisition functions exist (Frazier, 2018; Garnett, 2023; Shahriari et al., 2016; Wang et al., 2022) and are often applied in a batch setting (Bailey et al., 2024; Garnett, 2023; Hunt, 2020; Yang et al., 2025), where multiple sequences are proposed at once. This increases experimental efficiency through parallel experimental characterization, but requires batch diversity to maximize the knowledge gained. Current methods maximize the joint uncertainty of the batch by assuming normal distributed predictions (Bailey et al., 2024) or a heuristic selecting the top candidates according to a utility measure that does not account for interdependencies between candidates (Yang et al., 2025).

Following the appeal of riboswitches for information processing and the complexity of their design, the goal of this study is the engineering of digital like NAND hybrid riboswitches through a novel machine-learning framework for sequence design. Precise emulation of NAND behavior by the hybrid riboswitch is required to ensure its viability in a wide range of applications. Essential to the design of this dual-input RNA device is a method actively coping with the sensitivity of hybrid riboswitches with respect to single nucleotide exchanges and explicitly considering the switching behavior to achieve. Specifically, the requirement for fine-tuning the four different states of the device to achieve near digital NAND behavior raises needs beyond those required for classical dynamic range maximization.

To this end, we propose a multistep design approach incorporating *in vivo* screening and batch Bayesian optimization for score-driven sequence design. Our sequence design framework implements the batch acquisition function by using an ensemble neural network (Cao et al., 2020; Dietterich, 2000; Ganaie et al., 2022; Krogh and Vedelsby, 1994), which extends a custom pre-trained transformer encoder (Devlin et al., 2019; Vaswani et al., 2017) as surrogate and Kriging Believer (KB) (Desautels et al., 2014; Garnett, 2023; Ginsbourger et al., 2008; Hunt, 2020) as batching strategy. By considering batch diversity and allowing arbitrary distributions, this goes beyond recent works. In addition, the network quantifies the certainty associated with its predictions and is thus applicable to low throughput regimes such as those used to characterize hybrid riboswitches with high precision. Inspired by the design of hybrid riboswitches engineered for maximum dynamic range (Kelvin et al., 2025), we combined a tetracycline with a neomycin riboswitch and performed an *in vivo* screening to identify a variant showing NAND-like behavior, i.e. significant regulatory activity only in the presence of both ligands. This construct serves as input to our sequence design framework. The riboswitch design itself implements a design-build-test-learn (DBTL) cycle, in which the *in silico* candidate selection and the *in vivo* characterization alternate. The novel hybrid riboswitch we report here is capable of precisely emulating NAND logic through direct binding interactions, resulting in the stabilization of its compact structure only in the simultaneous presence of two different small molecule triggers. In addition, we present our versatile machine learning sequence design framework based on batch Bayesian optimization, which is adaptable to a wide range of design tasks, and provide insights into the method.

## Results

### Template Selection

In a first step we screened for a hybrid riboswitch that implements Boolean NAND-like behavior, exhibiting significant inhibition of gene expression only in the simultaneous presence of both ligands, while maintaining equally high expression levels in the absence of either or both ligands (Figure 1**a**).

**Figure 1:**
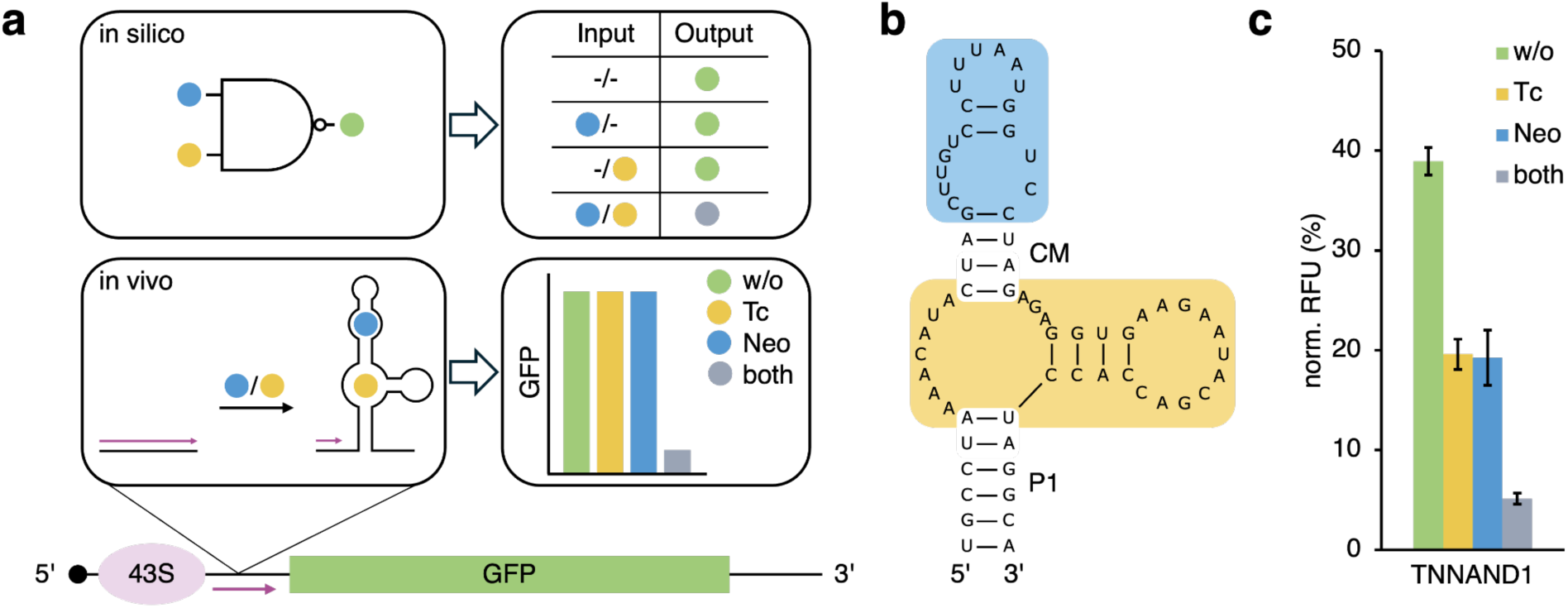
Translation regulation in yeast by a tetracycline-neomycin hybrid riboswitch following Boolean logic. (**a**) Schematic representation of the roadblock mechanism of translation regulation in yeast being performed by a tetracycline-neomycin hybrid riboswitch following NAND logic (blue: neomycin; yellow: tetracycline). (**b**) 2D structure prediction of the TNNAND1 hybrid riboswitch. The binding pockets for neomycin and tetracycline have been highlighted in blue and yellow, respectively. (**c**) Effect of TNNAND1 on GFP expression in the absence (w/o) and presence of 250 µM tetracycline (Tc), neomycin (Neo), or both. S. cerevisiae RS453 were transformed with the plasmid pcBB06 containing TNNAND1 in the 5’ UTR of a constitutively expressed gfp gene. Cells were incubated under the respective ligand conditions for 24 h. GFP values were normalized to constitutively expressed mCherry to eliminate cell to cell expression variability. pCBB06 without an insert was used as the blank and its values were subtracted from all other measurements. The fluorescence values of each measurement were normalized to their respective ligand-specific positive control (pcBB05). All measurements were performed in duplicates and repeated twice on separate days.

For the initial NAND screening we choose to combine two well-established riboswitches that recognize tetracycline (Berens et al., 2001) and neomycin (Weigand et al., 2008), respectively. Thereby, the P2 stem of the tetracycline riboswitch is being replaced by a truncated version of the neomycin riboswitch (Figure 1**b**). We refer to the stem sequence connecting the two binding pockets as a communication module. We designed the hybrid riboswitch to contain a short communication module that enforces close spatial proximity between the two binding pockets, thus allowing direct interactions. Additionally, it was shown that expression levels regulated by such a device can be modulated when using different versions of the closing P1 stem (Kelvin et al., 2025), an effect that was previously observed for the original tetracycline riboswitch (Groher et al., 2019) (Figure S1). We used the tetracycline-neomycin variant TN6 (Kelvin et al., 2025) for inspiration, as it exhibits only moderate dynamic ranges in the individual presence of either ligand. We considered this behavior to be a favorable trait for an initial NAND screening template. An additional UA base pair was added to the base of the P1 stem to increase overall stability of the construct. The sequence of the resulting construct TNNAND1 is depicted in Figure 1**b**. The effect of TNNAND1 on gene expression was analysed by inserting the construct into the 5‘ UTR of a *gfp* reporter gene directly upstream of the start codon. GFP expression was measured after a 24 h incubation period in the absence and presence of either or both ligands (w/o; 250 µM tetracycline (Tc); 250 µM neomycin (Neo); 250 µM both). The dynamic range achieved by TNNAND1 in the dual-input state is 7.6-fold, while remaining 2-fold in the presence of only one ligand (Figure 1**c**). While the deviation between the state without ligand and the individual ligand states means that TNNAND1 does not achieve any defined logic behavior, we found the similarity of the two individual ligand states to be a good starting point for further engineering.

### Screening for a NAND Communication Module

We designed a library of different riboswitch sequences through randomization of the TNNAND1 communication module between the binding pockets for tetracycline and neomycin, except for the conserved closing GC base pair of the internal bulge of the neomycin binding pocket, resulting in an N6 pool. For the initial *in vivo* screening we sorted yeast cells containing the different N6 variants based on their GFP fluorescence. However, sorting specifically for high fluorescence in the presence of one or no ligand and low fluorescence in the presence of both ligands was not sufficient to guarantee the discovery of candidates with NAND behavior. This is because all variants contained in the N6 pool have their own sequence-specific expression levels and candidates with the most NAND-like behavior are not guaranteed to also achieve high expression levels in the absence of both ligands or a high dynamic range in their presence, making a standard sorting approach difficult. To circumvent this issue, we divided the maximum GFP fluorescence range (0-100%) into subpopulations using equally sized gates, thereby creating a sorting grid that we could use for all ligand conditions. We were able to determine the exact sequences contained in each subpopulation using next-generation sequencing (NGS). Because we used the same grid for all conditions, subpopulations originating from a specific gate always contained candidates corresponding to its fluorescence range. This strategy allowed us to approximate the switching pattern of each sequence contained in the N6 pool by comparing its occurrence in different subpopulations for all ligand conditions.

We determined the maximum GFP fluorescence range using control plasmids and divided it into eight equally sized gates on a linear scale ranging from low to high fluorescence. Two additional outer gates cover the fluorescence ranges occupied by the controls. We then used a yeast library containing the N6 pool to inoculate four different cultures, one for each ligand condition (w/o; 250 µM Tc; 250 µM Neo; 250 µM both). Using the predetermined grid of gates, we divided each culture into subpopulations. We sorted 12,500 candidates per gate for each ligand condition. Plasmid DNA was extracted from each subpopulation and the contained sequences were verified using NGS. We then approximated the switching pattern of each sequence by comparing its occurrence in different subpopulations for every ligand condition, with subpopulations originating from gates in the lower fluorescence range corresponding to low expression levels and vice versa (Figure 2**a**).

**Figure 2:**
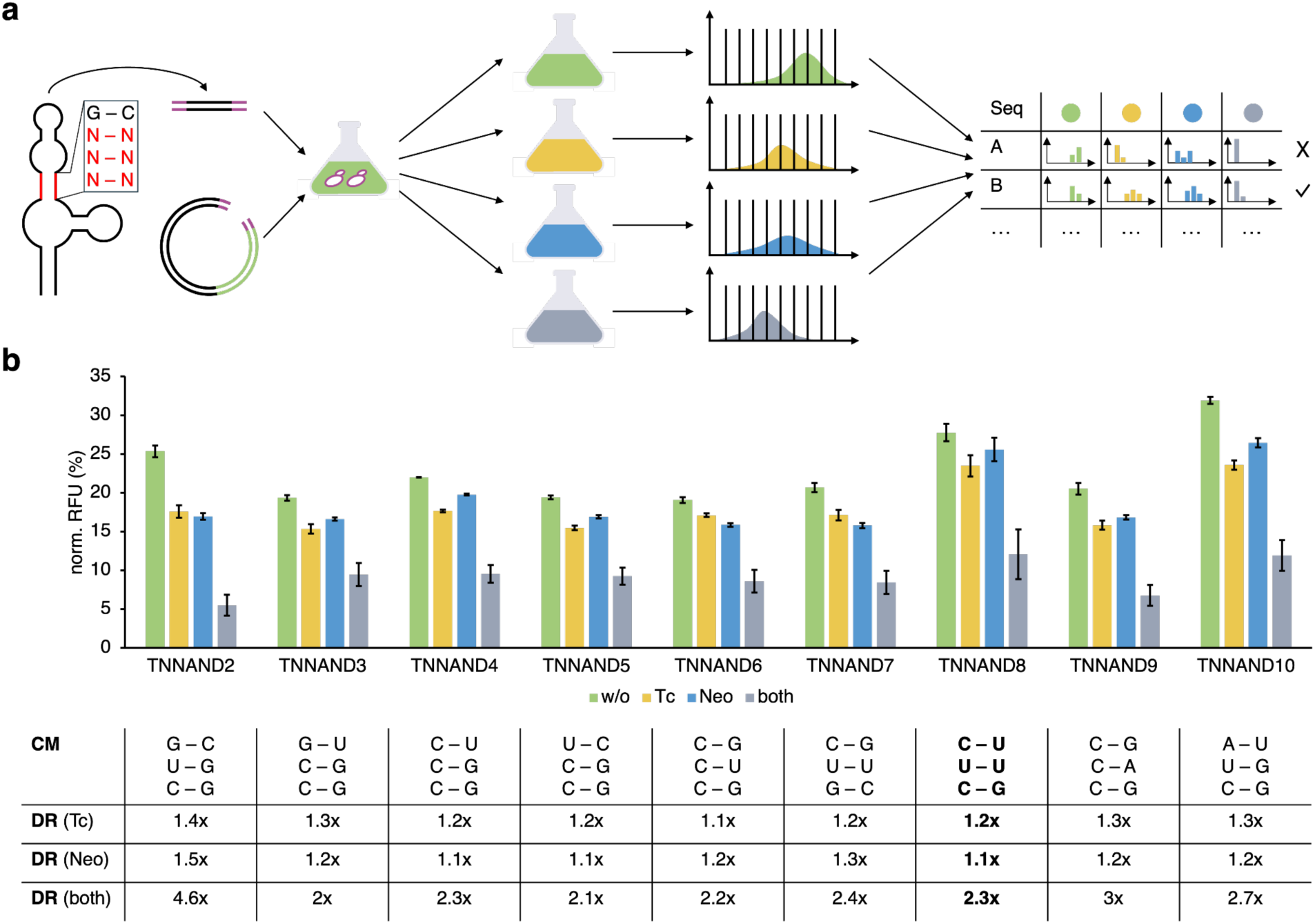
NAND communication module screening top candidates. (**a**) Screening workflow. A yeast library containing the TNNAND1 sequence with a randomized communication module sequence was created via homologous recombination. Cultures for each ligand condition were inoculated and sorted through the same grid of equally sized gates. The plasmid DNA from each subpopulation was extracted and analyzed via NGS. The resulting datasets were used to approximate the switching pattern of each sequence discovered under all four ligand conditions. (**b**) The influence on GFP expression of candidates that emulate NAND behavior was measured in the absence (w/o) and presence of 250 µM tetracycline (Tc), neomycin (Neo), or both. S. cerevisiae RS453 were transformed with the plasmid pcBB06 containing the respective candidate in the 5’ UTR of a constitutively expressed gfp gene. Cells were incubated under the respective ligand conditions for 24 h. GFP values were normalized to constitutively expressed mCherry to eliminate cell to cell expression variability. pCBB06 without an insert was used as the blank and its values were subtracted from all other measurements. The fluorescence values of each measurement were normalized to its respective ligand-specific positive control (pcBB05). All measurements were performed in duplicates and repeated twice on separate days. The communication module sequence of each candidate is depicted below its respective measurement results.

We identified sequences occurring in lower gates for the double ligand condition, while showing similar distributions across gates representing higher fluorescence for all other ligand conditions using the NGS datasets. We then made an expert selection of sequences that demonstrated the most promising NAND-like behavior based on their overall performance across all ligand conditions. We proceeded to clone the chosen candidates TNNAND2-10 and individually measure their effect on GFP expression. The communication module sequences of TNNAND2-10, as well as their dynamic ranges for each ligand condition are depicted in Figure 2**b**. All candidates present NAND logic to some degree, with dynamic ranges for the individual ligand conditions remaining low in comparison to the double ligand condition (e.g., TNNAND2: Tc = 1.4x; Neo = 1.5x; both = 4.6x). However, constructs with a higher dynamic range in the presence of both ligands also present a stronger decrease in GFP values for the individual ligand conditions. Constructs with a low dynamic range in the presence of either ligand suffer from poor regulation, even in the presence of both ligands (e.g., TNNAND8: Tc = 1.2x; Neo = 1.1x; both = 2.3x). We identified TNNAND8 as the most interesting candidate for further optimization. Given that the sequence of the tetracycline riboswitch closing stem P1 strongly influences switching behavior (Groher et al., 2019; Kelvin et al., 2025), our next step was to engineer a TNNAND8-specific P1 stem to improve dynamic range while maintaining the NAND behavior conferred by the new communication module.

Instead of six nucleotides (three base pair long communication module), which yield 4^6^ = 4096 combinations in total, the P1 stem region includes five or even more base pairs, increasing the combinatorial search space to over one million combinations. Due to this exponential increase in complexity, the characterization of expression levels by cloning and screening sequence pools of this size and variety becomes challenging. In particular, this requires the significant increase of the number of cells sorted per fluorescence gate to maximize the pool coverage and preserve the desired precision. However, the increase of time needed for sorting also poses challenges on the accuracy, as the measurements have to match the timings of the experimental protocols.

### Bayesian Optimization for Navigating the Sequence Space

The combinatorial increase in complexity renders purely experimental based sequence design inefficient and we thus next investigated the use of Bayesian optimization in combination with sequence-to-function surrogate models for efficient and functionality-driven riboswitch stem design. Incorporating individual construct measurements, the learned sequence-to-function model can assess the functionality of yet unexplored riboswitch sequences, while Bayesian optimization for sequence selection efficiently trades off exploration and exploitation to cope with the combinatorial complexity of the sequence space.

The P1 stem design is a particular instance of biological sequence design, i.e., the task of assigning nucleotides in an engineered construct to achieve a desired behavior. This behavior can be specific expression levels, dynamic range maximization, particular sequence features or - as in our case- the emulation of a particular Boolean function. We assess the degree of functionality via a scoring function *S*(⋅), which is based on experimental characterization. Consequently, we formalize the design task in terms of an optimization problem with the ideal design given by

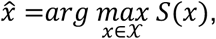

where 𝒳 is the sequence design space and *x*& maximizes the score *S*(⋅). To solve this design task, we build a new sequence design framework on the basis of Bayesian optimization.

Core to Bayesian optimization is the model’s belief 𝘱[*S*(*x*) | *D_j_*], expressed as a probability distribution over scores conditioned on the knowledge or data *D_j_* available at optimization round *j*. For a particular sequence *x* ∈ 𝒳, the belief represents a probabilistic sequence-to-function mapping giving rise to not only the expected function value in terms of *E*[*S*(*x*′) | *D_j_*] but also the associated uncertainty. This serves as a surrogate to guide Bayesian optimization also in regions not yet explored by experimental assessment. In particular, the acquisition function, which determines the candidates to evaluate next, selects the sequence *x* maximizing the utility function *u*(*x*). This utility function is derived from the probabilistic predictions of the surrogate model.

We select the upper confidence bound (UCB) (Cox and John, 1992; Desautels et al., 2014; Garnett, 2023; Hunt, 2020; Shahriari et al., 2016; Srinivas et al., 2012; Wang et al., 2022) as the acquisition function for the sequence design task because of interpretability, expressiveness, and simplicity of evaluation in combination with deep ensemble models. The UCB estimates the utility of each sequence by an upper bound on the achievable score. Formally, the utility function *u*(*x*) is defined to satisfy

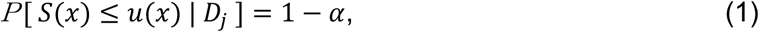

where *P*[*S*(*x*) ≤ *s*|*D_j_*] is the cumulative distribution function (CDF) corresponding to 𝘱[*S*(*x*)|*D_j_*] and *α* ∈ [0,1] defines the desired level of confidence and can be used to trade off exploitation (*α* → 0.5) and exploration (*α* → 0). For example, *α* = 0.05 leads to an utility *u*(*x*) such that, according to the model, the probability of *S*(*x*) exceeding *u*(*x*) is at most 5%. The ensemble model as surrogate allows us to define the UCB directly in terms of Equation (1) by using the quantiles of the empirical distribution, i.e.,

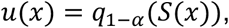

where *q*_1−*α*_(*S*(*x*)) refers to the 1 − *α* quantile function. The sequence *x*′ maximizing *u*(⋅) is then chosen to be evaluated experimentally (see Figure 3**a**).

**Figure 3:**
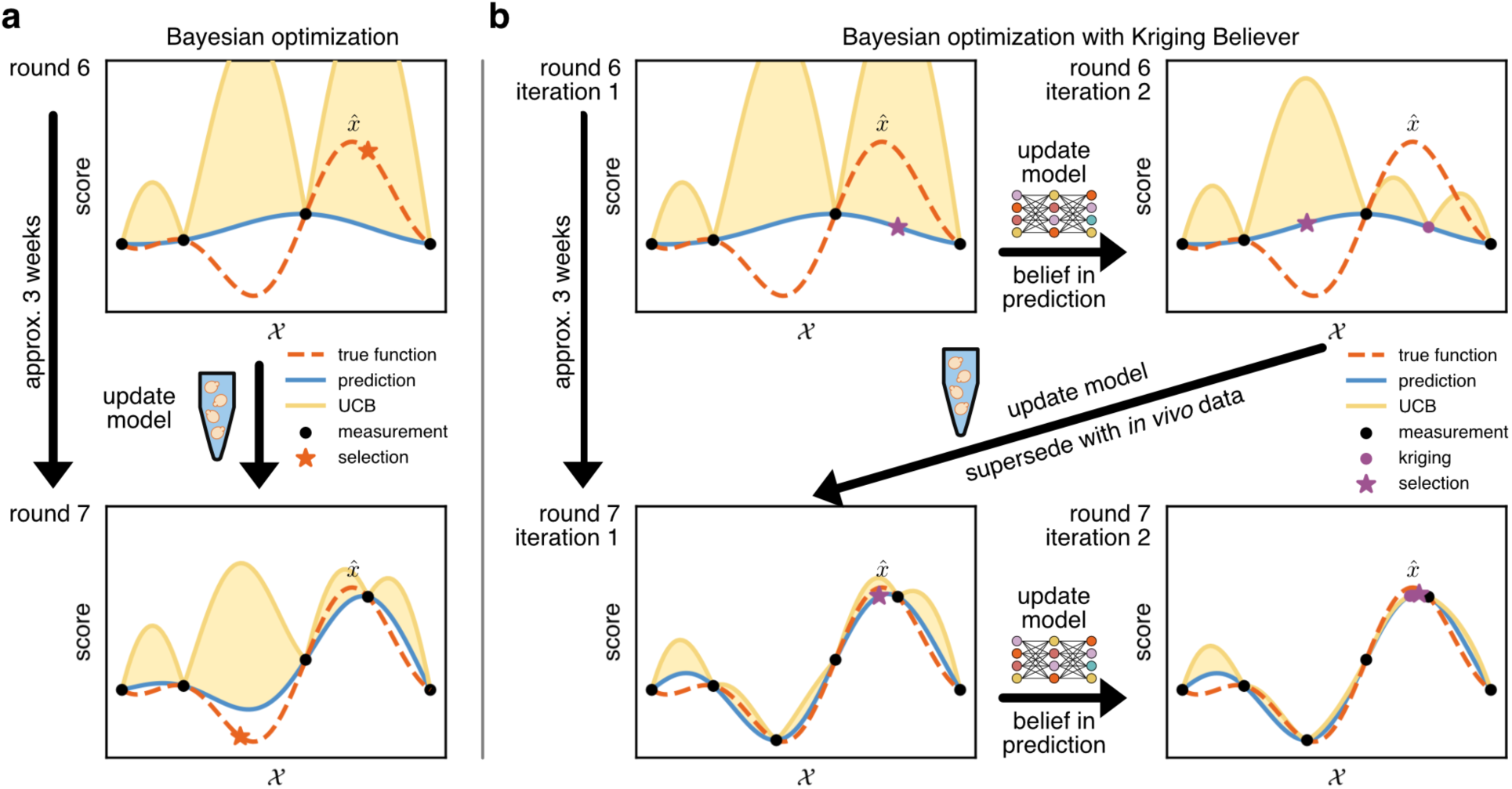
Batch Bayesian optimization. Side by side comparison of Bayesian optimization (panel **a**) and batch Bayesian optimization employing Kriging Believer (panel **b**). (**a)** Bayesian optimization proposes a single sequence (orange star) maximizing the utility function (here UCB, yellow line) per round. The surrogate model’s belief (blue line and UCB) is updated with the in vivo sequence characterization. (**b)** Kriging Believer introduces multiple iterations per batch Bayesian optimization round and also selects the sequence maximizing the utility function. Between iterations, the model’s belief is updated with the predicted value (purple star). Upon selection of all sequences for the current batch (in this example *m* = 2), the model’s belief is updated with the in vivo characterization of the batch. Errors introduced by kriging are superseded between rounds (compare round 6 iteration 2 and round 7 iteration 1).

#### Experimental Efficiency through Batch Creation

We devise a novel Bayesian optimization method for the batch setting (Desautels et al., 2014; Garnett, 2023; Hunt, 2020) to make use of the parallelism of the assay characterizing the riboswitches, where multiple candidate sequences *x*′_1_, …, *x*′_*m*_ need to be proposed at once. Repeatedly querying the acquisition function without updating the model’s belief will either (1) select the same riboswitch or, by excluding case (1) explicitly, (2) select sequences with high proximity in terms of the score *S*(⋅). The consequence of both cases is the absence of diversity, again being detrimental to experimental efficiency.

We achieve batch diversity by extending our proposal mechanism by the Kriging Believer (KB) technique (Desautels et al., 2014; Garnett, 2023; Ginsbourger et al., 2008; Hunt, 2020), explicitly excluding case (1). The idea of Kriging Believer is to skip the experimental evaluation and instead update the model’s belief with the predicted values. As such, the uncertainty vanishes for candidates in the current batch and decreases in their vicinity. Upon selecting the *m* candidates, the batch *x*′_1_, …, *x*′_*m*_ is experimentally characterized and the model is updated with the true measurements. Figure 3**b** visualizes this technique for *m* = 2, allowing a side-by-side comparison of Bayesian optimization and Kriging Believer based batch Bayesian optimization. For clarity, we refer to a round as a single run of batch Bayesian optimization and to an iteration as a single step of Bayesian optimization as part of a batch Bayesian optimization round (compare Figures 3**a** and 3**b**).

While Kriging Believer blends nicely with Bayesian optimization and provides an appealing, intuitive, and straightforward approach for the extension to the batch setting, the approach may raise concerns regarding error propagation. Every evaluation of the utility function -with the exception of the first iteration-explicitly depends on the predictions made in the previous iterations (see Figure 3**b** round 6, iteration2). These predictions are usually not exact and prediction errors can negatively affect future predictions. However, these errors do not propagate into the next round of batch Bayesian optimization, as all predictions are superseded by the experimental characterization (compare Figure 3**b** round 6, iteration 2 to round 7, iteration 1). Besides, future rounds of batch Bayesian optimization as well as the uncertainty reduction in the proximity of sequences in the batch are independent of the particular predictions (see Figure 3**b** round 6, iteration 1). As such, beneficial candidate sequences previously discarded because of incorrect predictions can be reconsidered later.

### Deep Ensemble Neural Network for Uncertainty quantification

We propose the use of an ensemble neural network (Cao et al., 2020; Dietterich, 2000; Ganaie et al., 2022; Krogh and Vedelsby, 1994) representing a probabilistic sequence-to-function mapping to guide the Bayesian optimization in the sequence space. By learning the sequence-to-function relations from experimental data, this model allows to assess yet uncharacterized sequences.

We create each of the *n* networks in the ensemble from a sequence encoder featuring a specifically designed BERT like transformer encoder (Devlin et al., 2019; Vaswani et al., 2017) and a fully connected regression network (MLP) (Russell and Norvig, 2021). To train the ensemble, we employ a two stage training process. First, we pre-train the sequence encoder on a task specific sequence dataset to learn the structure inherent to the sequence space under consideration. The masked language model (MLM) technique, which is widely used in natural language processing (NLP), learns in-sequence relationships by randomly masking parts of a sequence and requiring the model to predict the masked parts. As previous studies have shown the minor quality of embeddings learned by relying on MLM alone, we extend our pre-training with a novel metric learning task that we derived from the triplet margin loss used in classification tasks (Hoffer and Ailon, 2015; Malkiel et al., 2022).

As depicted in Figure 4**a**, we combine both objectives by forming a triplet network of identical sequence encoders. For a single training of the network, we first select sequence *x*_1_ and a corresponding hard negative *x*_2_ (see **Methods**). From these, we derive the anchor *x*_*a*_ = *x*_1_, the positive *x*_*p*_ = *mask*(*x*_1_) and the negative *x*_*n*_ = *mask*(*x*_2_) by masking, as described previously (Devlin et al., 2019) and outlined in **Methods**. The triplet network computes the corresponding embeddings *v*_*a*_, *v*_*p*_, *v*_*n*_ and predicts the masked parts of *x*_*p*_ and *x*_*n*_. Our overall training loss is then defined as

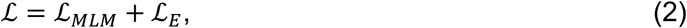

where ℒ_*MLM*_ is the masked language model loss (Devlin et al., 2019) and ℒ_*E*_ is the triplet margin loss (Hoffer and Ailon, 2015; Malkiel et al., 2022) with margin parameter *γ* and euclidean norm

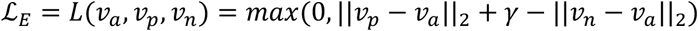

**Figure 4:**
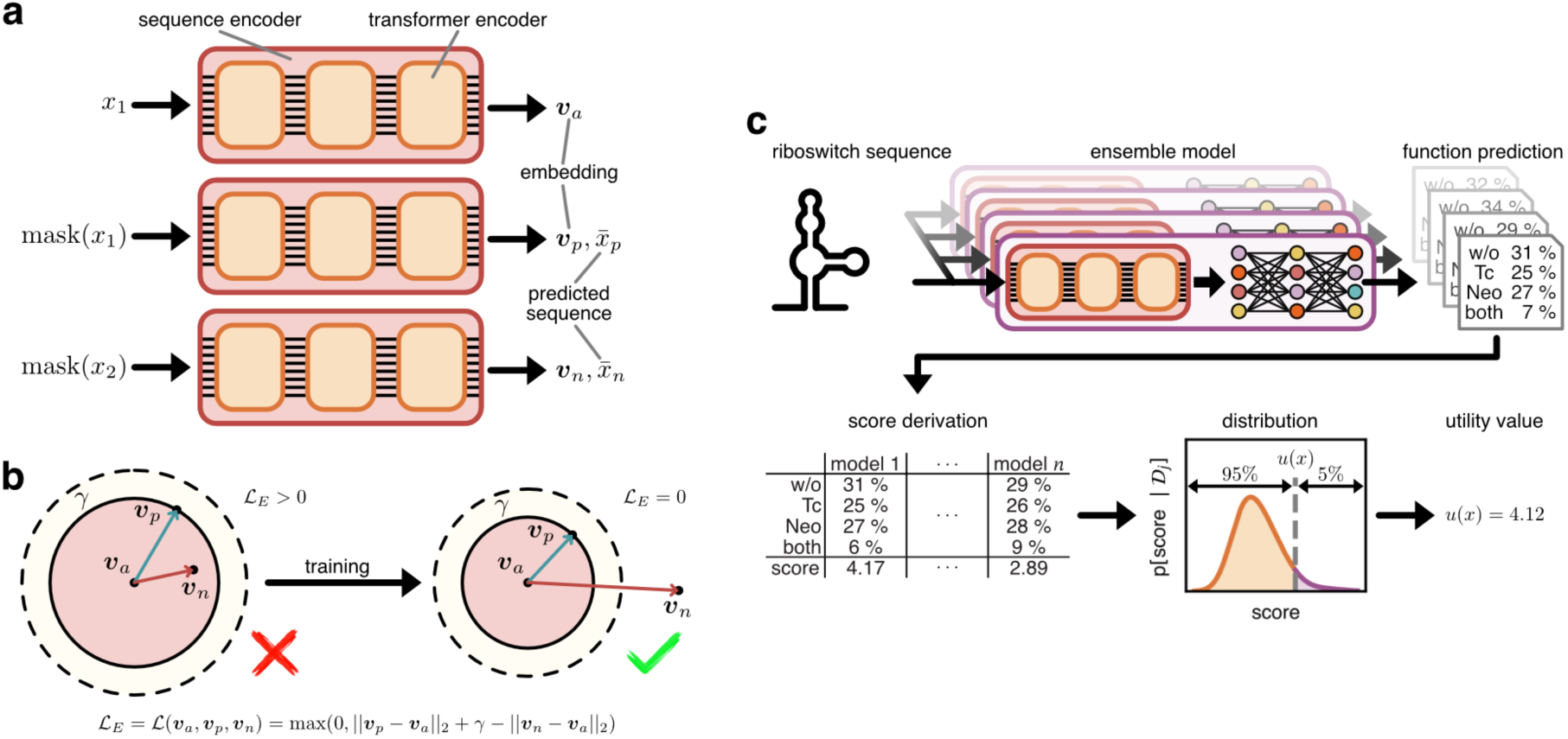
Pre-training and deep ensemble model. **(a)** Triplet network setup for pre-training of the sequence encoder. The same sequence encoder (here depicted three times) is applied to the sequences *x*_1_, the masked version of *x*_1_, and the masked version of *x*_2_. The output values are then considered by the loss function (Equation (2)). **(b)** Visualization of the triplet margin loss. Training maximizes the embedding distance between distinct sequences. **(c)** Sequence-to-function neural network ensemble as Bayesian optimization utility function instantiating the UCB.

Given the pre-trained sequence encoder, the second stage of training focuses on the sequence-to-function ensemble model. In particular, a fine-tuning approach is applied where each network in the ensemble is trained independently of each other on the same dataset. Despite the same training data, the random initialization of the parameters of the MLP ensures the diversity of the ensemble (Dietterich, 2000; Ganaie et al., 2022).

As a surrogate model for Bayesian optimization, the trained ensemble predicts the empirical distribution of the score from which the utility of the sequence is derived (Figure 4**c**), while the fine-tuning process is repeated from scratch for every Bayesian optimization iteration.

### DBTL Cycle for NAND P1 Stem Design

We next apply the batch Bayesian optimization framework we developed for the design of the TNNAND8 P1 stem by combining it with *in vivo* riboswitch screenings. This combination allows for a functionality-driven design of the P1 stem while using the individual construct measurements required for accurate design and simultaneously tackling the sequence space complexity. In particular, our method comprises three stages: (i) the design of a riboswitch sequence pool, (ii) the screening of the sequence pool for dataset creation, and (iii) our Bayesian optimization framework for the efficient and effective design of riboswitch sequences. In the initial step (Figure 5**a**), the sequence pool is created by using the sequence of TNNAND8 as the basis for the generation of a pool of partially randomized oligonucleotides (randomization of P1 stem). By employing fluorescence-activated cell sorting (Figure 5**b**), we enrich sequences that demonstrate sufficient dynamic range between the absence and presence of both ligands. Only these sequences are taken into account by Bayesian optimization. As depicted in Figure 5**c**, this instantiates a DBTL cycle. After multiple rounds, this approach results in riboswitch sequences maximizing the desired NAND functionality. We describe each of these steps below and provide a flowchart representation in Figure S2.

**Figure 5:**
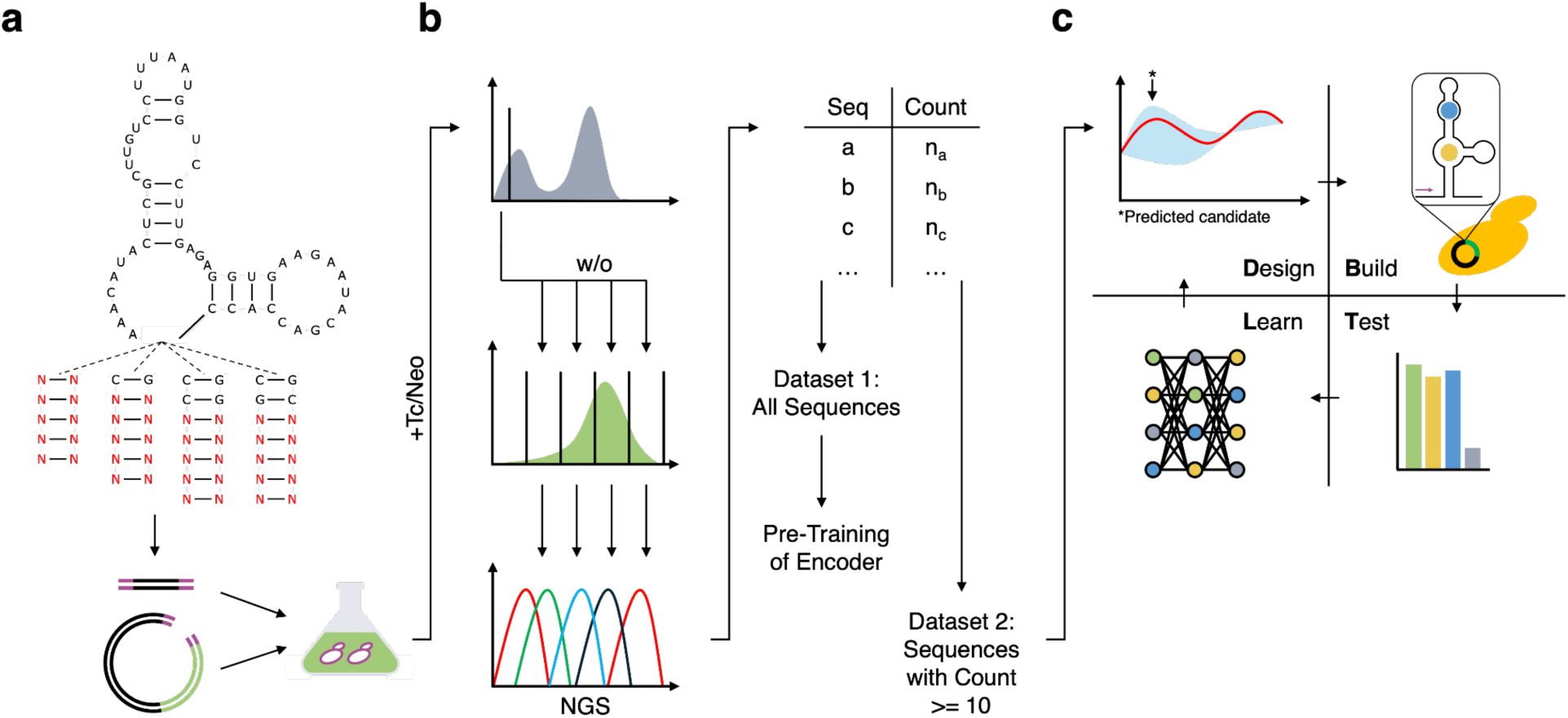
Overview on machine learning guided NAND hybrid riboswitch design. (**a**) TNNAND8 P1 stem sequence pool design. Yeast libraries were generated through homologous recombination with the linearized vector pCBB06. (**b**) Sequence pool and dataset creation. Candidates with high dynamic range in the presence of both ligands were enriched through cell sorting. Four different gates were used for the ligand-free state sorting of candidates previously enriched for low reporter expression in the presence of both ligands to gain a better understanding of riboswitch behavior at different dynamic ranges. NGS was used to determine the sequence space contained in each enriched subpopulation. For pre-training of the encoder a first dataset containing all detected sequence variants was used. A second dataset only containing sequences discovered ten times or more was used for model training. (**c**) DBTL cycle. Based on the initial training, the model predicted potentially interesting candidates for in vivo validation (Design). The respective constructs were cloned (Build), measured in yeast (Test), and the resulting data was used to train the model for the next iteration (Learn).

The screening process focuses on enriching riboswitches with a high dynamic range between presence and absence of both ligands. With the goal of increasing sequence diversity without losing pool coverage, we derive four distinct sequence pool designs of TNNAND8. Of note, P1 stems of the tetracycline aptamer conferring improved regulation are likely to contain certain closing base pair combinations below the binding pocket (C-G, C-G/C-G, C-G/G-C) (Groher et al., 2019). Therefore, each pool is designed to contain a five base pair long randomized sequence within the tetracycline P1 stem and either no additional closing base pair, or one of the aforementioned closing base pair combinations (Figure 5**a**, Pools T1-4 from left to right). We separately transformed *S. cerevisiae* cultures with the four resulting N10 pools (10^4^ = 1,048,576 variants; transformation efficiency per pool approx. 5 x 10^6^ cfu/µg plasmid). Afterwards, we combined the four cultures.

With the sorting range being defined on the basis of control plasmids, we first sorted the combined N10 pools for low fluorescence values (0-10%) in the presence of both ligands to remove riboswitches without function (1.2 million sorted cells) (Figure 5**b**). The sorted subpopulation was allowed to grow and regenerate before the next sorting step. For the second sorting we created equally sized gates dividing GFP values between 20-100% into four different subpopulations. We did this to differentiate between candidates with lower and higher dynamic ranges to avoid enriching only high performing hybrid riboswitches that have abolished NAND logic. We based the amount of sorted cells per gate on the size of the respective subpopulation to maximize coverage of all contained sequence variants (20-40% GFP: 360k events; 40-60% GFP: 36k events; 60-80% GFP: 10k events; 80-100% GFP: 10k events).

We again extracted and amplified the sequences to analyze them via NGS. In preparation for the computational framework, we created two datasets. Dataset 1 contains all sequences identified via NGS and will serve as a dataset for the pre-training of the sequence-to-function model. In contrast to this, dataset 2 will serve as the candidate sequence pool from which the batch Bayesian optimization selects the sequences for experimental characterization. To increase the experimental efficiency by removing sequencing artefacts, we required a read count of at least 10 for sequences to be part of dataset 2.

### Application of Bayesian Optimization to Riboswitch Design

We define the scoring function *S*(⋅) to be the worst-case dynamic range with respect to NAND behavior to assess the riboswitch performance in a manner accessible to the sequence design. It assesses the dynamic range between the smallest expression level representing an Boolean ON state (ligand conditions w/o, Tc, and Neo) and the one representing the Boolean OFF state (ligand condition both). Formally, this score is defined as

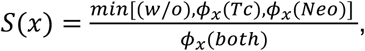

where 𝜙_*x*_(⋅) is the expression level of sequence *x* for some ligand condition.

For the sequence design framework, we use a batch size of *m* = 16 sequences, an ensemble size of *n* = 100 models, and configure the acquisition function UCB to bound 95 % of predicted scores by setting *α* = 0.05. For pre-training the sequence-encoder, dataset 1 consists of 36,925 riboswitch sequences. The candidate sequence pool, from which we select the sequences during our optimization, is dataset 2 and has a size of 3,256 sequences (see Figure 6 for sequence space visualization).

**Figure 6:**
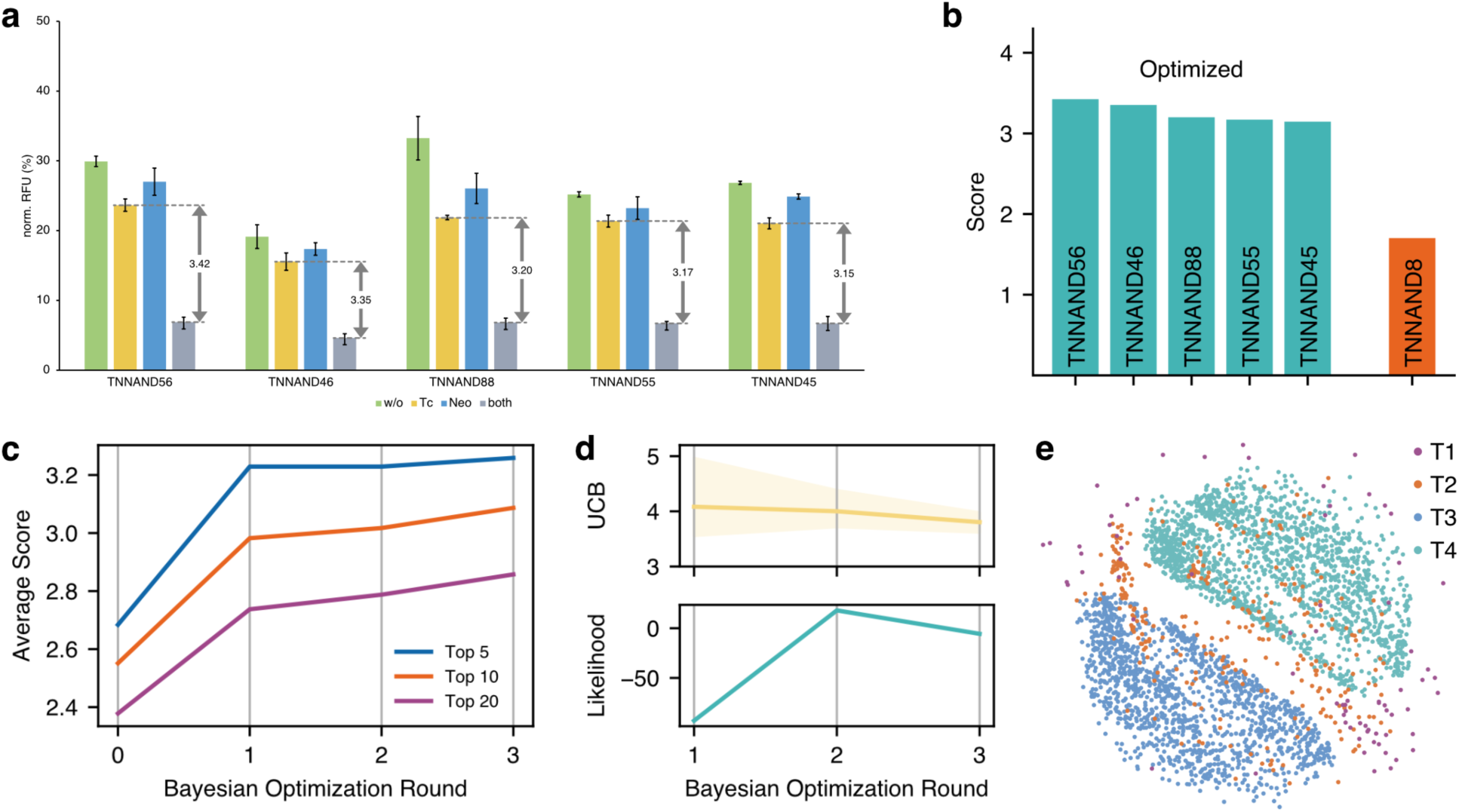
NAND riboswitch design results. **(a)** Relative fluorescence of the top 5 NAND riboswitches TNNAND56 (score 3.42), TNNAND46 (score 3.35), TNNAND88 (score 3.20), TNNAND55 (score 3.17), and TNNAND45 (score 3.15), ordered by descending performance quantified by the worst-case fold-change score. **(b)** Worst-case fold-change scores of top 5 NAND riboswitches compared to the initial construct TNNAND8 (score 1.70). TNNAND56, as the best performing NAND riboswitch, achieves a 2-fold improvement. **(c-e)** Evolution of quantities over the Bayesian optimization rounds. **(c)** The average score of the top 5, top 10, and top 20 sequences in each round of Bayesian optimization. **(d)** Per round statistics of upper confidence bound (UCB) (upper panel) and predictive log-likelihood (lower panel). For the UCB, the average (solid line) and minimum to maximum interval (shaded area) over the 16 sequences per round are presented. The descending trend indicates the decrease of model uncertainty. The Log- likelihood of measurement values is obtained by applying a normal closure to the ensemble prediction. Increase in log-likelihood indicates the improvement in model performance through the increased knowledge. **(e)** Sequence space visualization by multidimensional scaling (MDS) of pairwise Levenshtein distances between sequences. Clustering of sequences corresponding to templates T3 and T4 is likely due to these templates being the most prevalent ones.

For the selection of the 34 initial riboswitches to characterize (TNNAND11-45, TNNAND29 is not included)^1^, we employed a semi-random approach. Focusing on potential function and sequence space coverage, we rationally selected the majority of riboswitches and supplemented them with candidates that cover most sequences not yet considered. For this purpose, we use sequence similarity on the basis of the Levenshtein distance. All chosen candidates, despite not being sorted for a specific logic gate pattern, retained NAND logic gate behavior (Figure S3). To us, these results clearly indicate that the TNNAND8 communication module specifically enforces NAND logic gate behavior.

Starting the first round of our Bayesian optimization technique, the ensemble regression model is trained on the 34 sequences and the corresponding expression levels. For all sequences in the sequence pool, we evaluate the upper confidence bound and select the sequence with the highest utility value. The next iterations of this round continue by augmenting the current knowledge with the estimated expression levels of the selected sequence (the Kriging Believer step), training the regression model, and selecting the sequence with the highest upper confidence bound. Upon selection of all sequences for the current batch, we continue with the *in vivo* evaluation, the build and test steps of the DBTL cycle in Figure 5**c**.

In this first round of batch Bayesian optimization (TNNAND46-61) we found 11 riboswitch sequences that performed better than the previous average and three that outperformed the previous best. Among those 16 sequences, there are also four sequences without NAND functionality (Figure S4). Noticeably, their stems have at most two canonical base pairs. Guided by those insights and the rationale of the riboswitch mechanism, we consider the absence of base pairs in the stem causal for the absence of any switching behavior. Therefore, we removed sequences featuring less than 3 base pairs (including G-U pairs) in the stem from the sequence pool, to focus exploration on promising candidates. This reduces the size of the sequence pool to 3,177 sequences.

Including the expression levels of the newly evaluated sequences into our dataset extends it to 50 entries. Rounds two and three of our Bayesian optimization proposed 32 further sequences. However, the *in vivo* characterization revealed that these rounds did not yield a new overall best NAND riboswitch. In addition, the predicted utilities of yet unknown riboswitches decreased, wherefore we terminated optimization with only 82 riboswitches being evaluated *in vivo*.

Considering the performance of the newly designed riboswitch sequences, Figure 6**a** presents for each inducer combination the average expression levels of the five best NAND riboswitches. Noticeably, the expression levels representing Boolean ON states (i.e. for input combinations w/o, Tc, and Neo) are well separated from the ones representing OFF states (i.e. for input combination both). In addition, the lowest ON state is on average only 22.1 % below the highest, with TNNAND88 having the highest deviation of 34.2 %. In combination with the average worst-case dynamic range of 3.26 among our top five, these riboswitches implement near digital NAND logic *in vivo*. Setting the performance of the top five in relation to the initial construct TNNAND8 (see Figure 6**b**), they all outperform TNNAND8 (score 1.7) score wise, while our method improved the worst-case dynamic range by 2 fold for the best candidate TNNAND56 (score 3.42).

Investigating the convergence of the proposed technique, Figure 6**c** highlights that the presented method continuously improves the average scores of the top 5, top 10, and top 20 riboswitches with each round of Bayesian optimization. While the 16 sequences selected in round 1 led to the largest improvements, the increases in later rounds are mostly of incremental nature. However, especially with respect to the top 10 and the top 20, the increase of performance between round 1 and round 3 is still significant and sustainably enriches the abundance of well performing NAND riboswitches. Taking the UCB values of the proposed riboswitches into account (see Figure 6**d**), we observe a decrease in network uncertainty and utility attributed to yet unknown sequences. In view of the performance of the best riboswitch TNNAND56, featuring a score of 3.42, and the increase of measurement likelihood (see lower panel **d**) being evidence of improved model accuracy, the UCB values confirm the reasonability of terminating optimization after three rounds of batch Bayesian optimization.

### Analysis of the Bayesian Optimizer

In this work, Bayesian optimization is central for the design of riboswitches. Therefore, we here investigate relevant quantities for Bayesian optimization in order to provide further insights into the selection process. Figure 7 visualizes the model’s view of the sequence space at the beginning of each round (round four corresponding to panel **d** is not executed). Starting with panel **a** corresponding to the first round, the distance between UCB (representing the 0.95 % quantile) and mean estimate is evident for the large uncertainty within the model. Reconsidering the measurement likelihood in Figure 6**d**, this estimate of uncertainty correlates well with the predictive performance of the model. Considering the models knowledge at the beginning of round 1, we see that the characterized riboswitches cover the whole performance range from almost non-functional to NAND behavior, while model uncertainty vanishes at the corresponding positions (downward spikes in UCB). As in general the UCB values are high, the riboswitches selected are distributed over the whole sequence space, including candidates with low estimated performance but high uncertainty, being exemplary for exploration, and ones with lower uncertainty but higher performance estimates, being exemplary for exploitation. Following the definition of the acquisition function, the selected candidate sequences should correspond to peaks in the UCB utility function. However, this does not match our observations in the panels. The reason for this is, that the presented UCB is exemplary for the models knowledge when starting the Bayesian optimization round. This is not identical to the model’s belief in the iteration the candidate riboswitch is selected. Causal to this is the Kriging Believer approach, which updates the model’s belief in each iteration to reflect the knowledge gained through the incorporation of the previously selected riboswitches.

**Figure 7:**
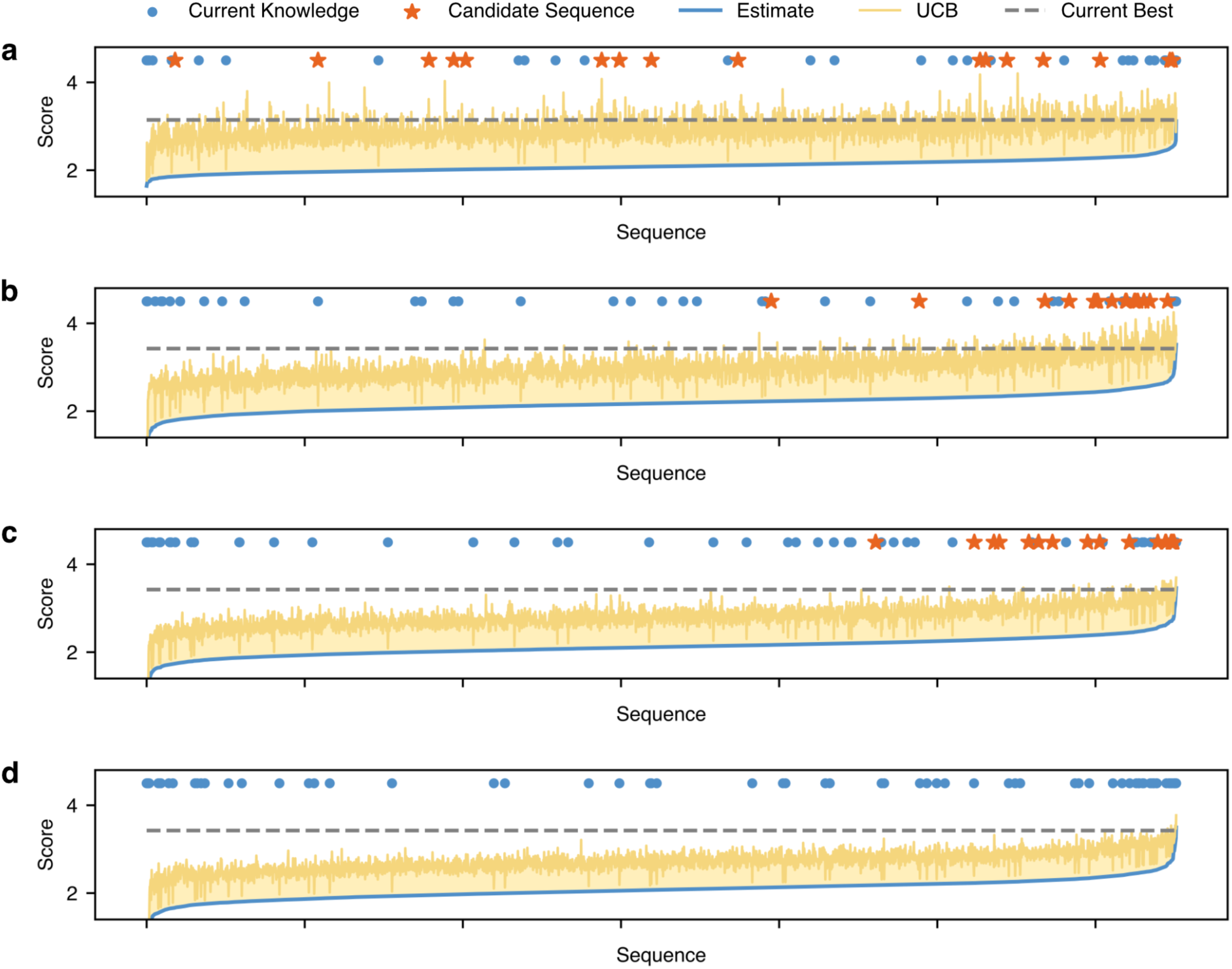
Batch Bayesian optimization rounds. Visualizations of the models predictions *μ*(*x*) (blue line) and the corresponding upper confidence bound *u*(*x*) (UCB, yellow area) for each sequence in the candidate sequence pool on the basis of the knowledge at the start of round one (**a**), two (**b**), and three (**a**), with panel **(d)** representing the state for an unexecuted fourth round. In addition, markers indicate current knowledge (blue points) and proposed sequences to evaluate (orange stars). Sequences are ordered per round by ascending predicted score. Deviations between UCB peaks and selected candidate sequences result from different model states for visualization and sequence selection.

Continuing with round 2 (see Figure 7**b**), we observe a general drop in uncertainty (deviation between UCB and mean) and a preference for riboswitches with higher estimated performance in comparison to round 1 (panel **a** for reference). Indeed, the information gained through the experimental data corresponding to the riboswitches selected in round 1 refines the predictions (see Figure 6**d**), also in the realm of low performing sequences. As such, also for these riboswitches uncertainty decreases, reducing the probability of their selection due to the combination of low estimated performance and medium uncertainty levels. On the other side of the performance spectrum, uncertainty is causal for candidate selection, as the difference in the expected performance among predicted top performers decreases. With respect to the diversity of selected riboswitches, the model’s knowledge did lead to preferring exploitation, with the exception of some candidates being chosen for the purpose of exploration.

Round 3 manifests the trend of decreasing uncertainty and the selection of candidates with higher estimated performance (see Figure 7**c**). Especially in the range of riboswitches with medium estimated score, single peaks in uncertainty are causal for the selection of a riboswitch. However, we also do notice that in most cases the distance between selected candidates is larger than in round 2. This indicates the success of the Kriging Believer approach in reducing uncertainty in the proximity of riboswitches already part of the current batch. Besides, the selected riboswitches again indicate the trade off between exploration and exploitation by considering sequences with diverse estimated performance. This is in contrast to greedy optimization, which focuses only on the sequences with the highest estimated performance.

After three rounds of batch Bayesian optimization, the model’s uncertainty decreased significantly (see Figure 7**d**) and the upper confidence bound is in general below the already identified best scoring riboswitch. This indicates the vanishing chance of identifying riboswitches with superior performance. The scores of the evaluated riboswitches span from 0.91, representing an unfunctional riboswitch, to 3.42 in the case of TNNAND56. From the 48 riboswitch sequences selected by batch Bayesian optimization only 14 have a score below 1.70 (score of TNNAND8), while 15 switches exceed this score by at least 50 %.

### Sequence Characteristics of NAND Hybrid Riboswitches

On the basis of the expression profiles of the 82 riboswitch sequences collected during optimization, we here investigate the relation between sequence and function. Focusing on the best performing riboswitch first, Figure 8**a** gives rise to the nucleotide frequency per stem position for the top 10 sequences according to the NAND score. While for most positions each nucleotide appears at least once, the preference for templates T3 (5 out of 10) and T4 (4 out of 10) in the top 10 is evident at position 7, as only a single sequence corresponds to template T2 (1 out of 10), featuring a stem of length six and templates T3 and T4 do not divert at this position. At position 6, nucleotides G and C are almost equally frequent, again reflecting the fraction of templates in the top 10. Considering template T1, featuring the shortest stem of length five without any predefined base pairs, this template is not present in the top 10 and only three -in general non functional-corresponding sequences have been evaluated in total. Positions 1 to 5 are not predetermined by the templates and present a more heterogeneous choice of nucleotides. In general, nucleotide G is most prominent on the right side, with its complementary C occurring slightly less often on the left.

**Figure 8:**
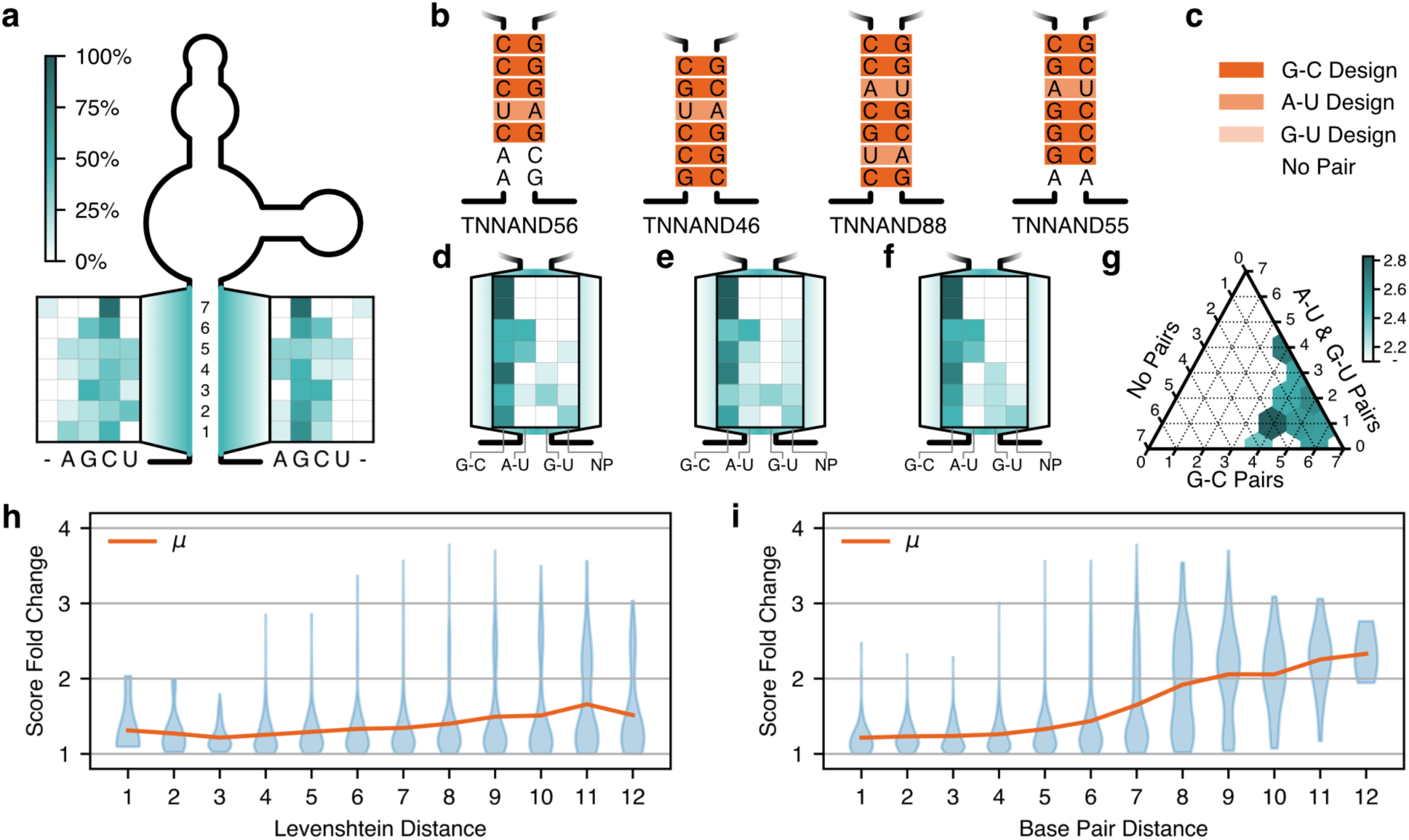
Sequence insights into riboswitch design. **(a)** Relative frequency of nucleotides of top 10 riboswitch sequences, where - indicates the absence of a base through a shorter stem. **(b)** Sequences of top 4 riboswitches designed in this work with highlighting of base pairs following the legend in **c**. **(d-f)** Relative frequency of base pairs in top 10 of the riboswitches with respect to the score (**d**), high expression in the “w/o” case (**e**), and low expression in the “both” case (**f**). **(g)** Average scores of top 40 riboswitches in dependence to the number of base pairs, where we differentiate between no base pairing (NP), G-C base pairing (G-C pairs), and A-U or G-U base pairing (A-U & G-U Pairs). **(h)** Riboswitch sequence robustness analysis on the basis of the distribution of the fold change in score (lower is better) in dependence to the Levenshtein distance between sequences, with substitutions, insertions, and deletions weighted equally. Small score fold change means almost identical scores. **i:** The distribution of the fold change in score in relation to a base pair distance, where the distance between two distinct base pairs is 1 and between a base pair and no base pair is 2.

Narrowing down to the top 4 riboswitches, their actual sequences are displayed in Figure 8**b**, with position wise base pairs highlighted. Noticeably, the best performing riboswitch TNNAND56 features only five base pairs, all being in the upper part, while the others feature up to seven. In addition, there is a strong prevalence for G-C base pairs (19 in total) over A-U base pairs (5 in total). Here, TNNAND56 again diverts from the others in featuring three G-C base pairs in the upper part of the stem. Extending our considerations to the top 10 performers with respect to the score, Figure 8**d** highlights that G-C base pairs are the most prevalent ones, with A-U pairs occurring only at positions 2, 4, and 5. The non-canonical G-U pair and the absence of base pairs is mainly present at the lower part of the stem. Figures 8**e** and **f** continue this evaluation by focusing on the 10 riboswitches with highest expression levels in the “w/o” case (panel **e**) and the 10 with lowest expression levels in the “both” case (panel **f**). Especially by comparing these two, one observes that different base pairing patterns seem optimal. The increased absence of base pairs indicates that high expression in the “w/o” case is best achieved by weakening the base pairing. Contrary to that, the strong G-C base pairs are most frequently used at each position when considering the sequences with the least leakiness in the “both” case.

Reconsidering the score, Figure 8**g** visualizes the average score in dependence to the number of base pairs per type for the 40 best NAND riboswitches. All riboswitches have at most three positions without base pairing, while on average, the best performing configurations have four pairs of type G-C and one of either type A-U or G-U. Besides those, riboswitches with seven base pairs perform better than average.

An important aspect for sequence design and the predictability of functionality is the robustness of the sequence with respect to altering single nucleotides. To investigate this, we present in Figure 8**h** the fold change in score -which we use as performance similarity metric-between sequences with defined Levenshtein distance (the fold change in score is *s*_1_/*s*_2_ if *s*_1_ ≥ *s*_2_ else *s*_2_/*s*_1_, with *s*_1_ = *S*(*x*_1_) and *s*_2_ = *S*(*x*_2_)). Focusing on the mean first, we observe a slight increase in score fold change for increasing Levenshtein distance. However, additionally taking the distribution information into account, we observe that distributions for different Levenshtein distances are almost similar, while already a Levenshtein distance of 1 can lead to a 2 fold change in performance. As by intuition and guidance of the other panels in Figure 8 the number of base pairs and their type is an important quantity, we additionally introduce a Base pair distance. The base pair distance weights the distance between two distinct base pairs with 1 and between no base pair and a base pair with 2. We present the corresponding results in Figure 8**i**, with the mean presenting an almost monotonic behavior and spanning a much larger range in comparison to panel **h**. In addition, the distributions differ more significantly, indicating an improved categorization of sequences.

## Discussion

RNA-based regulators such as synthetic riboswitches offer enormous potential for the design of complex genetic circuits. Despite this, riboswitches are difficult to engineer due to their sensitivity to sequence changes, with even single nucleotide exchanges having a major impact on regulated expression levels. Because sequence-function relationships are often unpredictable, riboswitch design in many cases remains dependent on time-consuming empirical methods. Machine learning approaches have attempted to bridge this gap by combining high-throughput wet lab data generation with sequence-to-function surrogate models. However, these studies have focused exclusively on the generation of single-input RNA devices with maximized dynamic range (Angenent-Mari et al., 2020; Groher et al., 2019; Valeri et al., 2020). Our hybrid riboswitch architecture surpasses these devices in complexity by allowing the use of two different inputs to simultaneously influence the switching behavior, thereby enabling the implementation of Boolean logic. This complexity is further enhanced by the fact that hybrid riboswitch function is based solely on structure stabilization induced by small-molecule triggers, unlike other riboswitch architectures that rely on strand displacement to facilitate regulation (Angenent-Mari et al., 2020; Valeri et al., 2020). Until now, the design of hybrid riboswitches has focused solely on maximizing dynamic range to create highly efficient regulators that take advantage of the additional stability provided by a second binding pocket (Kelvin et al., 2025). However, since dynamic range is always dependent on the system being controlled, we have focused here on the logic behavior enabled by the dual input, with the specific goal of achieving near-digital NAND logic behavior.

Most dual-input riboswitch architectures attempting to implement Boolean logic rely on separate insertion of two different devices into the same mRNA. While these tandem riboswitches can be created from the use of two different constructs utilizing the same mechanism of regulation, like the roadblock mechanism in S. cerevisiae (Schneider et al., 2017; Boussebayle et al., 2019), simultaneous use of different regulation mechanisms is also conceivable (Spöring et al., 2020; Weigand and Suess, 2007). Tandem riboswitches following Boolean logic have not only been engineered synthetically, but have also been discovered in nature (Welz and Breaker, 2007). For example, a tandem construct containing aptamers for guanine and 5-phosphoribosyl-1-pyrophosphate (PRPP) discovered in bacteria was shown to achieve IMPLY logic by influencing a shared expression platform (terminator stem) through both binding pockets (Sherlock et al., 2018). The fact that no natural riboswitch implementing NAND logic has been discovered to date is viewed as an indicator that interactions between both binding pockets are a requirement for dual-input riboswitch architectures to achieve complex Boolean functions (Sherlock et al., 2022). Previously published synthetic riboswitches implementing NAND behavior circumvent this by using a self-cleaving ribozyme, a catalytically active RNA sequence, as a shared expression platform for two different aptamer domains (Win and Smolke, 2008; Felletti et al., 2016). These designs make use of the modular nature of ribozymes by replacing two stem-loop structures essential to their function with aptamer domains that stabilize only in the bound state, thus allowing ribozyme activity only in ligand presence. However, the spatial separation between both binding pockets means that no direct interaction is utilized for NAND behavior, as each aptamer domain separately functions as a NOT gate. In comparison, our design utilizes direct interactions between both binding pockets to facilitate logic behavior, thus encoding NAND logic directly into a single RNA structure without the need for a separate expression platform.

Besides classical design through means like directed evolution (Tuerk and Gold, 1990; Ellington and Szostak, 1990; Nutiu and Li, 2005; Stoltenburg et al., 2012; Boussebayle et al., 2019), machine learning based approaches for the design of regulatory RNA devices have been proposed (Angenent-Mari et al., 2020; Iwano et al., 2022; Valeri et al., 2020; Wong et al., 2024). A collaborative work (Angenent-Mari et al., 2020; Valeri et al., 2020) combines high throughput cell sorting and sequencing with deep learning based sequence-to-function models to predict RNA toehold-switch function. The presented STORM optimization pipeline uses the trained model to optimize the input RNA sequence via gradient ascent with respect to target expression values in the bound and unbound state. In comparison, our method employs batch Bayesian optimization and probabilistic sequence-to-function modeling, additionally giving rise to the predictive uncertainty. Both are essential in data scarce scenarios. Also a generative approach to RNA aptamers on the basis of structure-to-sequence models has been proposed (Wong et al., 2024). This approach was exemplified by generating structurally similar but sequence wise dissimilar variants of light-up aptamers. Our framework directly focuses on the functional behavior of the riboswitch and provides means to maximize the performance with respect to a functionality score.

With respect to Bayesian optimization, our method goes beyond recent works by combining a large ensemble neural network with a batch acquisition function employing Kriging Believer and the upper confidence bound criterion. In particular, previous works use greedy approaches, stochastic acquisition functions or approximate joint uncertainty maximization for batch generation (Yang et al., 2025; Bailey et al., 2024). To generate probabilistic predictions, they use significantly smaller ensembles (Yang et al., 2025) or techniques for a single neural network such as Laplace approximation, Monte Carlo dropout or Gaussian process heads (Bailey et al., 2024; Stanton et al., 2022). In comparison, our method explicitly conditions the selection of riboswitch sequences during batch generation on previously selected sequences via Kriging Believer to improve batch diversity. In addition, we are able to account for arbitrary empirical distributions through the large ensemble and the quantile-based upper confidence bound, thus overcoming any distributional assumption.

Using our method, we achieved the design of a tetracycline-neomycin hybrid riboswitch that accurately emulates NAND logic, showing relevant downregulation of reporter expression only - and exclusively - in the presence of both ligands. Using the deep batch Bayesian optimization, we were able to double the performance of the optimized candidate TNNAND56 in comparison to the initially discovered construct TNNAND8. By having a total sample size of only 82 candidates validated *in vivo*, our framework shows robust performance in this low throughput regime. The novel combination of batch Bayesian optimization on the basis of Kriging Believer with a deep ensemble neural network as surrogate sequence-to-function model shows strong performance, with the pre-training being a beneficial priming strategy. Despite the computational complexity increasing through ensembling and Kriging Believer, the duration of *in vivo* characterization significantly exceeds the time for *in silico* batch creation.

Future evolutions can integrate further knowledge into the modeling, for example by taking structural information into account (He et al., 2024; Kagaya et al., 2025; Lorenz et al., 2011; Mortimer et al., 2014; Shen et al., 2024; Wayment-Steele et al., 2022). In the context of Bayesian optimization, the integration of means to trade-off data quality and quantity (Frazier, 2018; Garnett, 2023) is an orthogonal aspect to the previous and opens up a new dimension, which is promising for its application to RNA design but also to other sequence design tasks. Here, it is important to emphasize that the method presented can be adapted to other constructs beyond NAND hybrid riboswitches. By adjusting the in vivo screenings and defining the sequence constraints and scoring objective for the *in silico* process, our framework can be extended to general 5’ UTR design and adapted to other sequence design tasks for biological engineering. As such, our method can complement experiment driven approaches by allowing fine grained adaptation of functionality through model-based and score-driven design, including constructs sensitive to single nucleotide changes.

## Methods

### Experimental Methods

#### Plasmid Assembly

All constructs were cloned using the vector pCBB06, a variant of the plasmid pCBB05 containing no *gfp* start codon, as previously described (Boussebayle et al., 2019). Backbone linearization was performed immediately upstream of the gfp gene using the restriction sites for AgeI-HF and NheI-HF (NEB). Insert sequences containing a riboswitch sequence and start codon were generated through hybridization of oligonucleotides (Table S1) with overhangs corresponding to the aforementioned restriction enzymes. Constructs were assembled through ligation (T4 DNA Ligase (NEB)) and used for the transformation of *Escherichia coli* TOP10 (Invitrogen) following a standard 42°C heat shock protocol. The PureYield^TM^ Plasmid Miniprep System (Promega) was used for plasmid extraction, with sequences being verified via Sanger Sequencing (Microsynth Seqlab).

#### Yeast Cultivation and GFP Measurements

*Saccharomyces cerevisiae* RS453α cells (MATα *ade*2-1 *trp*1-1 *can*1-100 *leu*2-3 *his*3-1 *ura*3-52) were transformed using the Frozen-EZ Yeast Transformation II Kit (Zymo Research). The cells were used for the inoculation of a 24-well plate, with each well (2x for each construct) containing 1.5 ml SCD-ura [0.2% YNB w/o AA (Difco), 0.55% ammonium sulphate (Roth), 2% glucose (Roth), 12 μg/ml adenine (SIGMA), 1× MEM amino acids (SIGMA), 2% agar (Oxoid)]. After a 24 h incubation period at 30°C and 450 rpm, a 1:1000 dilution in fresh media containing either no ligand, 250 µM tetracycline or neomycin, or 250 µM of both ligands was performed in a new 24-well plate. After another 24 h of incubation the fluorescence of GFP and mCherry was measured for 25,000 cells per construct (20 µl culture + 180 µl SCD-Ura in a 96-well plate) using the CytoFlex S cytometer (Beckman Coulter) (488 nm laser (GFP); 561 nm laser (mCherry)). Emission light was bandpass filtered at 510/520 nm and 610/620 nm. Cells containing the control plasmids (positive: pCBB05; negative: pCBB06) were treated equally to the constructs and measured in parallel. The GFP fluorescence of each candidate and control was normalized to its respective mCherry fluorescence and the values of the negative control were subtracted from all candidates and the positive control. The adjusted value of each sample was normalized to the positive control incubated with the equivalent ligand condition. A biological duplicate was measured for every candidate and every experiment was performed in technical triplicates at independent days.

#### Yeast Library Pool Generation

Hybrid riboswitch DNA pools containing randomized regions (Table S2) were amplified (Q5 High-Fidelity DNA polymerase (NEB)) to add regions homologous to the insertion site on the plasmid pCBB06. *S. cerevisiae* RS453α cells were transformed according to a high-efficiency electroporation protocol (Benatuil et al., 2010), using 1.5 µg linearized plasmid pCBB06 (AgeI-HF and NheI-HF) and 1.5 µg amplified pool DNA, as previously described ((Kelvin et al., 2025)). Prior to transformation, pool and plasmid DNA were purified using the Wizard SV Gel and PCR Clean-Up System (Promega). Two transformations were performed for each pool to increase transformation efficiency and the transformed cells were combined in 250 ml SCD-Ura in a 2 l culture flask and incubated for 48 h at 30°C and 110 rpm. After the initial 48 h incubation period, serial dilution steps of the liquid culture in 250 ml SCD-Ura were performed according to the electroporation protocol to avoid the occurrence of double-transformed cells (1/10, 1/30, 1/100; 24 h incubation after each dilution).

#### NAND Communication Module Screening

A yeast library culture (see above; sequence pool TNNAND1_N6) was used to inoculate 4x 250 ml SCD-Ura cultures (1:100; 2 l flasks). Each inoculated culture contained a different ligand condition (w/o; 250 µM tetracycline; 250 µM neomycin; 250 µM both). Positive (pCBB05) and negative (pCBB05) controls were generated by transforming *S. cerevisiae* RS453 cells with the respective plasmids using the Frozen-EZ Yeast Transformation II Kit (Zymo Research). Cultures were incubated at 30 °C for 24 h in a humidified incubator (shaking). Prior to sorting cells were diluted (1:10) in 1x PBS. For fluorescence-activated cell sorting the SONY SH800 cell sorter was used (487 nm laser (GFP), bandpass filter: 525/550 nm). Doublets (connected cells) and cells of irregular shape or size were excluded. The GFP fluorescence range (0-100%) for cell sorting was defined as the distance between the peaks of the controls. The fluorescence range was divided into eight equally sized gates on a linear scale, with two additional gates covering the fluorescence ranges occupied by the controls. For each ligand condition, 12,500 candidates per gate were sorted into a separate well (500 µl SCD-Ura; 1 ml SCD-Ura added after sorting) on a 24-well plate. The plates were sealed (Greiner Bio-One Breathseal membrane) and incubated for 48 h at 30 °C using a plate incubator. Afterwards, plasmid DNA was extracted from all wells using the Zymoprep^TM^ Yeast Plasmid Miniprep II kit (Zymo Research) according to the manufacturer’s instructions. The sequence region containing the partially randomized hybrid riboswitch was amplified and the PCR product was purified using the Wizard® SV Gel and PCR Clean-Up System (Promega) according to the manufacturer’s instructions. The sequence content of all samples was analyzed using NGS (AMPLICON-EZ, GENEWIZ). As the gates span the whole fluorescence range and do not overlap, we approximate each sequence’s fluorescence by the frequency weighted average of the fluorescence intensities assigned to each gate. This is done for all four ligand conditions, whereby the gate’s fluorescence intensity is defined as the average fluorescence of events in a gate.

#### NAND Hybrid Riboswitch Dynamic Range Screening

Hybrid riboswitch yeast libraries for the sequence pools TNNAND8_T1(N10)/T2(N10)/T3(N10)/T4(N10) were created (see Yeast Library Pool Generation) and combined into one culture (1 ml of each library added to one 100 ml SCD-Ura culture). Positive (pCBB05) and negative (pCBB05) controls were generated as previously described (see NAND Communication Module Screening). All cultures were incubated for 24 h at 30 °C in a humidified incubator (shaking). The combined library and the controls were each used to inoculate (1:100) two 10 ml SCD-Ura cultures, containing either no ligand or 250 µM tetracycline and neomycin, and again incubated for 24 h at 30 °C in a humidified incubator (shaking). Prior to sorting cells were diluted (1:10) in 10 ml fresh SCD-Ura. For fluorescence-activated cell sorting the CytoFlex SRT cell sorter (Beckman Coulter) was used (488 nm laser (GFP); 561 nm laser (mCherry); bandpass filter: 525/540 nm and 610/620 nm). Doublets (connected cells), cells of irregular shape or size and cells lacking constitutive mCherry expression were excluded. GFP fluorescence was normalized to mCherry fluorescence and the fluorescence range (0-100%) for cell sorting was defined as the distance between the peaks of the controls. In the first sorting step 10% of all cells exhibited low fluorescence (0-10%) in the presence of both ligands. 1.2 million cells were sorted into 5 ml fresh SCD-Ura medium. The sorted cells were incubated for 48 h at 30 °C in a humidified incubator (shaking). Afterwards, two new 10 ml cultures, one for each ligand condition (w/o; 250 µM tetracycline and neomycin), were inoculated (1:100) and incubated for 24 h as previously described. The high fluorescence area was divided into four gates (20-40%, 40-60%, 60-80%, 80-100%) for the second sorting step, in case NAND behavior would begin to be compromised at higher expression levels. For each gate the amount of sorted cells was chosen based on the percentage of the population it contained to guarantee complete coverage (20-40% GFP: 360k cells; 40-60% GFP: 36k cells; 60-80% GFP: 10k cells; 80-100% GFP: 10k cells). Cells were sorted into four separate 5 ml SCD-Ura cultures and incubated for 24 h as previously described. Plasmid DNA was extracted from all four cultures as previously described (see NAND Communication Module Screening). The sequence content of all samples was analyzed using NGS (AMPLICON-EZ, GENEWIZ).

### Machine Learning Methods

#### Bayesian Optimization

Bayesian optimization is a global optimization method using a surrogate model, in our case the deep ensemble neural network, to guide optimization. The surrogate model describes the belief 𝘱[*S*(*x*)|*D_j_*], the distribution over *S*(*x*) conditioned on the knowledge *D_j_* in the current round *j*. This knowledge is

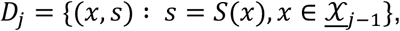

where <u>𝒳</u>*_j_*_−1_ is the set of sequences experimentally characterized in the previous rounds.

Derived from the model’s belief, the acquisition function assigns a utility value *u*(*x*) to each sequence *x* and selects the candidate by solving

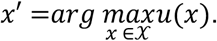

This optimization problem can be solved by means exhaustive enumeration, stochastic optimization or gradient based approaches exploiting differentiability of *u*(⋅). Enabled by the compact size of the final design space considered here, we solve this by exhaustive enumeration of the candidate sequence pool.

#### Kriging Believer

Kriging Believer is a natural extension of Bayesian optimization to the batch setting, as it replaces costly function evaluation by the model’s predictions and otherwise sticks to the sequential decision process. For iteration *i* of batch Bayesian optimization round *j*, the knowledge is

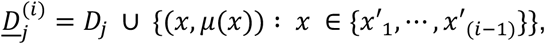

which is the union of *D_j_*, the actual measurements gathered at the start of round *j*, and the augmented dataset given by the predictions *μ*(*x*′_*k*_) = *E*[*S*(*x*′_*k*_)|<u>*D*</u>*_j_*^(*k*)^] for the sequences {*x*′_1_, ⋯, *x*′_(*i*−1)_} already part of the batch. By training the ensemble model from scratch (except the pre-trained encoder), the model’s belief is updated and explicitly conditions the predictions on the augmented values (i.e. 𝘱[*S*(*x*)|<u>*D*</u>^(*i*)^]). After batch creation (sequences *x*′_1_, ⋯, *x*′_*m*_), the knowledge for the next round of batch Bayesian optimization is

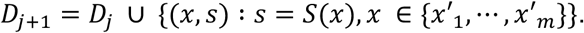

#### Encoder Model - Architecture and Training

The encoder model follows a BERT like Transformer encoder architecture (Devlin et al., 2019; Vaswani et al., 2017) including Multi-Head Attention (Vaswani et al., 2017). For tokenization, we build upon the 3-mer tokenizer of DNABERT (Ji et al., 2021) representing the construct sequence by 3-mer tokens such as [AGC]. Besides, we use [CLS] as the start token, [SEP] as the end token, and [PAD] as padding token. The combination of [CLS], [SEP], and [PAD] tokens allows to have inputs of varying length.

The encoder implementation consists of an embedding layer, positional encoding and six transformer encoder blocks. The embedding layer turns tokens into vectors of dimension *d*_*model*_ = 128, while we employ sinusoidal positional encoding (Vaswani et al., 2017). Each transformer encoder includes a multi-head self-attention mechanism consisting of 4 heads with dimension *d*_*attention*_ = 64. In comparison to the original work of Vaswani et al. (Vaswani et al., 2017) where *d*_*attention*_ = *d*_*model*_/ℎ for reasons of computational cost, we increase the attention dimension by a factor of two due to the smaller embedding dimension *d*_*model*_. The two layer position-wise feed-forward neural network, which is applied to each token position, has input and output dimension *d*_*model*_ = 128 and hidden dimension 384. Following Devlin et al. [Devlin et al. 2019], the first layer employs the Gaussian Error Linear Units (GELU) and the second a linear activation function. Both layers feature a dropout probability of 10%. The remainder of the transformer encoder follows the design of Vaswani et al. (Vaswani et al., 2017).

Our encoder model is trained on 36,925 unique riboswitch sequences obtained from sequencing. We apply train (70%), validation (15%) and test set (15%) splits. The considered riboswitch constructs have sequence lengths of up to 70 bases. After applying the 3-mer tokenization, the maximum length is 68 tokens. Taking start and end tokens into account, the maximum token sequence length we encounter is 70, while all token sequences shorter than that are extended with the [PAD] token. The training was carried out on a consumer grade NVIDIA GPU with a batch size of 64. Following four warmup epochs, the learning rate was determined by an exponential learning rate schedule with an initial learning rate of 8 ⋅ 10^−4^ and *γ* = 0.99. The learning rate is updated after each epoch and early stopping stops training after multiple iterations without improvements. This pre-training uses the loss defined by Equation (2) and includes the masked language model approach (Devlin et al., 2019) and the triplet margin loss (Schroff et al., 2015).

For the masked language model objective, we employ online masking (Liu et al., 2019) and otherwise follow the prescriptions of Devlin et al. 2019 (Devlin et al., 2019). In particular, 15% of the tokens in the token sequence (the special tokens [CLS], [SEP], and [PAD] are not considered) are selected randomly to be masked. From the tokens to mask, 80% will be replaced by the [MASK] token, 10% by a random 3-mer token, and 10% are not altered. For the masked token prediction, the encoder is extended by a position-wise single layer feed-forward network predicting the tokens at the respective positions. The masked language model loss is then the cross entropy loss of the predictions and the actual tokens, whereby only the masked positions are taken into account.

The triplet margin loss (Schroff et al., 2015) focuses on the quality of the embedding and requires an anchor, a positive and a negative. We here use the output of the [CLS] token as sequence embedding (yields embedding vectors of size *d*_*model*_ = 128). As anchors, we use the unmasked token sequences in the batch. The positive is obtained by applying masking to the anchor and the negative is the masked version of a token sequence identified via hard negative selection (Schroff et al., 2015). This selection criterion computes the embedding distance between each anchor in the batch and all other sequences and selects the one with the smallest non-zero distance. We then apply the triplet margin loss with a margin of *γ* = 0.98 to the embeddings derived by the sequence encoder.

The training loss is calculated as the superposition of the masked language model loss and the triplet margin loss. The best parameter configuration is selected based on the evaluation on the validation dataset and the final model achieved an accuracy of 96.9% on the test dataset.

#### Ensemble and Regression Model - Architecture and Training

The regression model is a fully connected multi-layer perceptron using the sequence embeddings as input and outputting the expression levels corresponding to the four riboswitch conformations. It consists of nine intermediate layers, each of dimension 256, and a linear output layer. The GELU function is used as the activation function and no dropout is used. To improve training performance, we use layer normalization after every three hidden layers.

Joining the pre-trained sequence encoder and the regression model yields the sequence-to-function neural network. *n* = 100 instantiations of this network constitute the ensemble model. To achieve diversity among models in the ensemble, the multi-layer-perceptrons are initialized randomly and the training procedure is designed to prevent ensemble collapse. In particular, dropout is only used within the encoder, as initial evaluations lead showed that further use of dropout negatively affects the network’s ability to differentiate known and unknown sequences. As hyperparameters we used a batch size of 4, 120 epochs, a learning rate of 10^−4^, a learning rate decay of 0.99, four initial warmup epochs, and a train validation split of 75 % and 25 %. The training includes the training of the sequence encoder. Each model in the ensemble is trained individually on the same dataset.

As loss function for the regression task we use the Kullback-Leibler divergence (KL divergence) (Kullback and Leibler, 1951). In particular, we consider the individual expression levels predicted -which are in the range between 0 and 1- as Bernoulli probabilities, characterizing the distribution of riboswitches expressing proteins or not. This bernoulli KL divergence for two Bernoulli distributions characterized by the probabilities p (the predicted value) and *q* (the target value) is given by

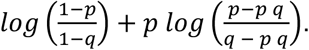

The KL divergence measures the -non metric- distance between two distributions and as such is minimal in case of identical distributions.

## Code and Data Availability

Code and data is available at https://github.com/Self-Organizing-Systems-TU-Darmstadt/NANDRiboswitchDesignByBayesOpt.

## Acknowledgments

D.K. and B.S. were supported by the Deutsche Forschungsgemeinschaft (DFG) CRC 902(A2). E.K. and H.K. were supported by ERC-PoC grant PLATE (101082333). Views and opinions expressed are however those of the author(s) only and do not necessarily reflect those of the funding agencies.

## Author Contributions

D.K. devised and executed wet lab experiments. E.K. devised and implemented the machine learning framework. D.K. and E.K. performed data analysis jointly. B.S. and H.K. provided funding. D.K., E.K., H.K., B.S. conceptualized the work and wrote the publication together.

## Competing Interests Statement

The authors declare no competing interests.

## Footnotes

1 TNNAND29 has the same sequence as TNNAND45 and was therefore not considered further.

## Notes

https://github.com/Self-Organizing-Systems-TU-Darmstadt/NANDRiboswitchDesignByBayesOpt

### Competing Interest Statement

The authors have declared no competing interest.

