## Supplementary Information for "NAND Hybrid Riboswitch Design by Deep Batch Bayesian Optimization"

DK, BS: Department of Biology, TU Darmstadt, Darmstadt, 64287, Germany
DK, EK, HK: Department of Electrical Engineering and Information Technology, TU Darmstadt, Darmstadt, 64283, Germany
DK, EK, HK, BS: Centre for Synthetic Biology, TU Darmstadt, Darmstadt, 64283, Germany
$ These authors contributed equally.

### Figure S1 - Tetracycline Riboswitch P1 Stem Variants

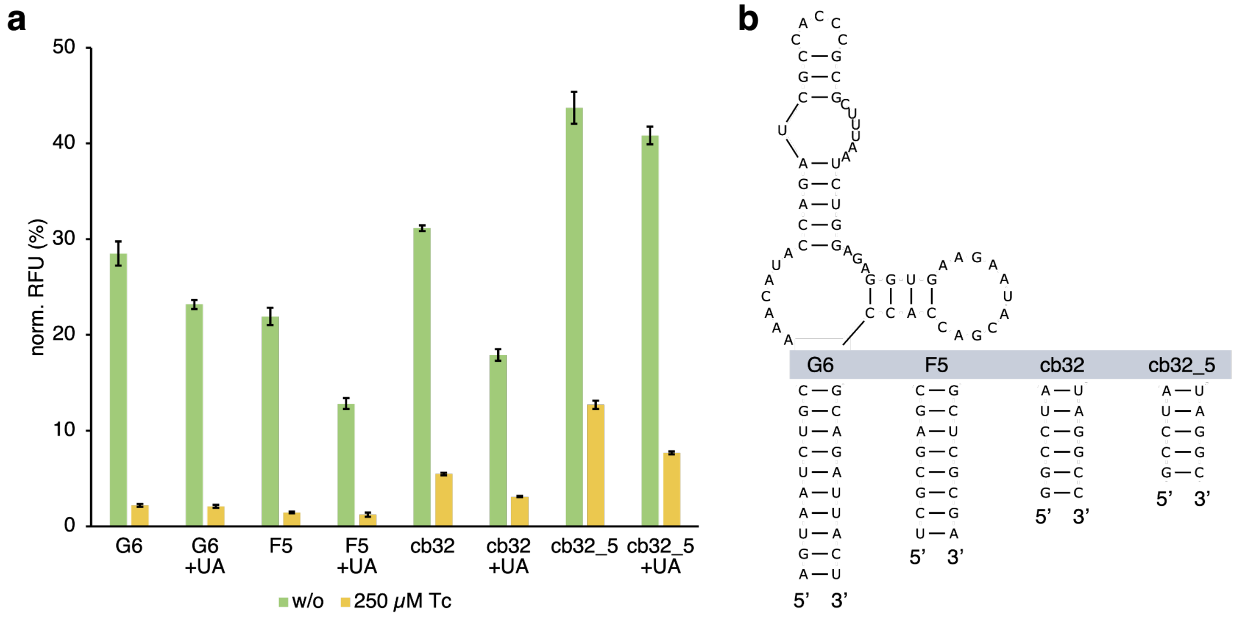

*Figure S1:* ***Tetracycline riboswitch P1 stem variants.*** *(**a**) The influence on GFP expression of each P1 stem variant was measured in the absence (w/o) and presence of 250 µM tetracycline (Tc). S. cerevisiae RS453 were transformed with the plasmid pcBB06 containing the respective candidate in the 5’ UTR of a constitutively expressed gfp gene. Cells were incubated under the respective ligand conditions for 24 h. GFP values were normalized to constitutively expressed mCherry to eliminate cell to cell expression variability. pCBB06 without an insert was used as the blank and its values were subtracted from all other measurements. The fluorescence values of each measurement were normalized to its respective ligand-specific positive control (pcBB05). All measurements were performed in duplicates and repeated twice on separate days. (**b**) 2D sequence representation of the Tc Riboswitch, including the different P1 stem variants.*

### Figure S2 - Flowchart of Deep Batch Bayesian Optimization

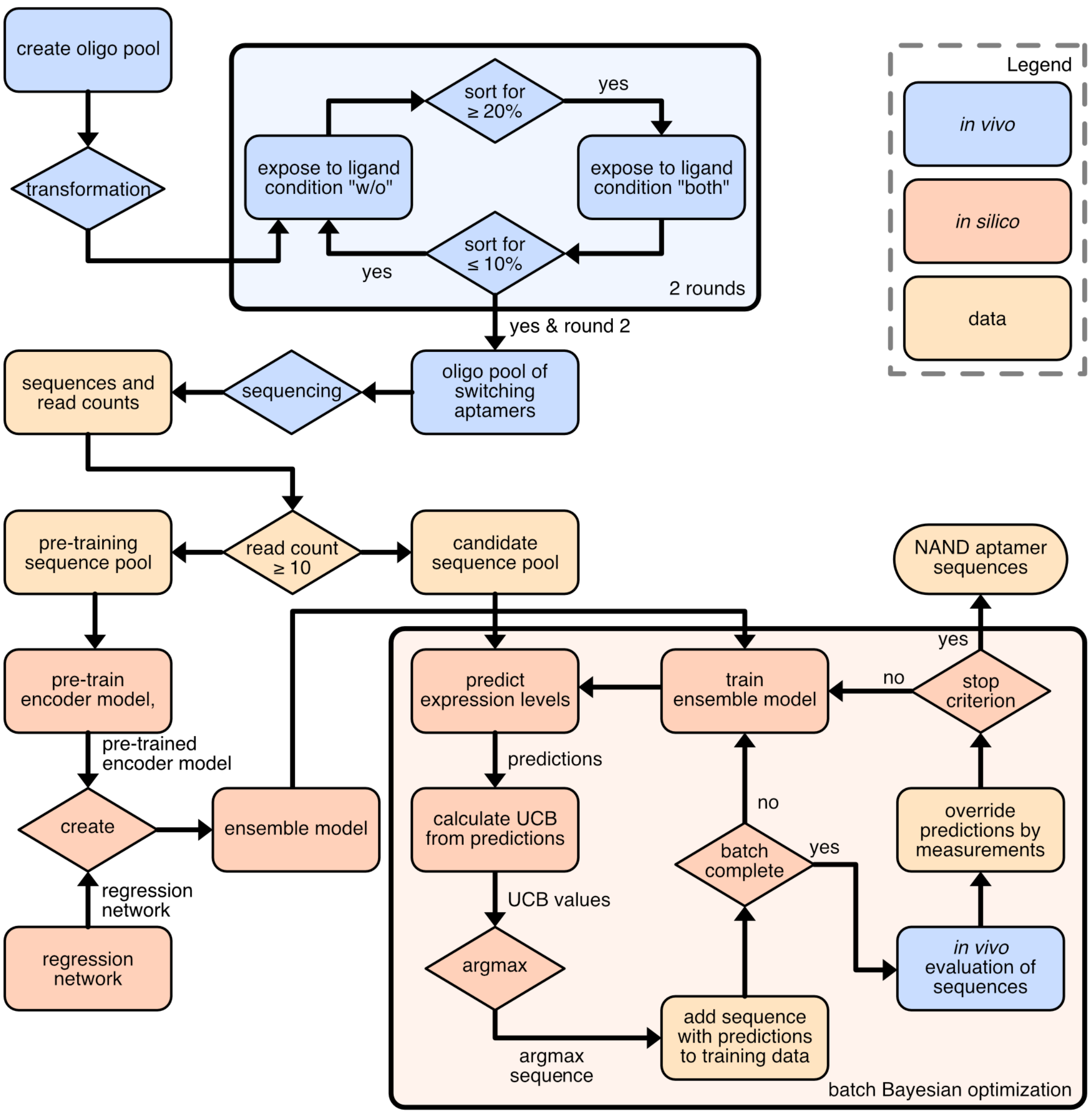

*Figure S2:* ***Deep Batch Bayesian Optimization Flowchart****. Starting at the initial oligo pool creation top right, this flowchart visualizes the process followed for the design of the hybrid riboswitch stem. The correspondence of different steps to either in vivo tasks, in silico tasks or data is highlighted by color and follows the legend given top right.*

### Figure S3 - ML Training Datasets (TNNAND8 P1 Variants)

**a**

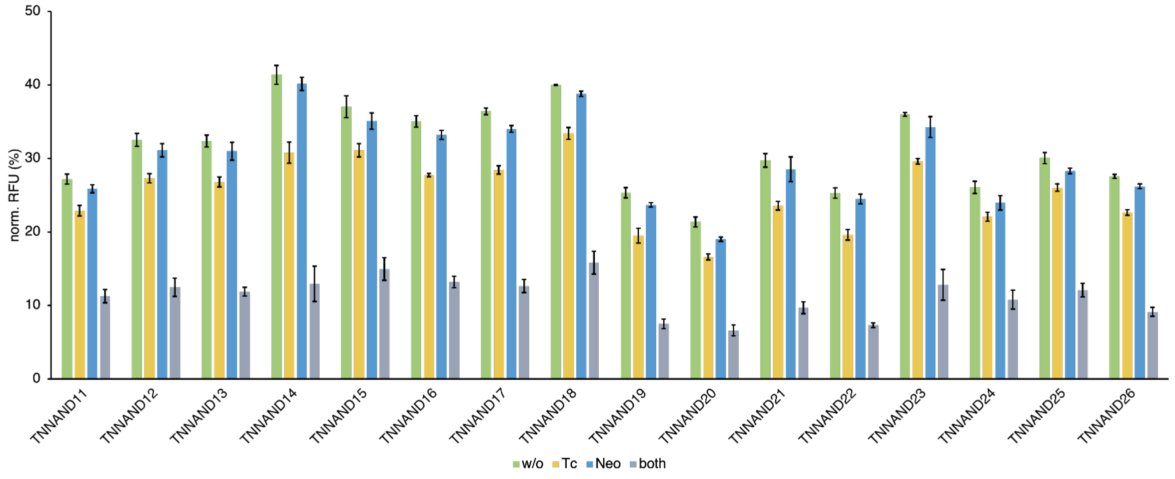

| | | | | | | | | | | | | | | | | |
|---|---|---|---|---|---|---|---|---|---|---|---|---|---|---|---|---|
| **DR** (Tc) | 1.2x | 1.2x | 1.2x | 1.3x | 1.2x | 1.3x | 1.3x | 1.2x | 1.3x | 1.3x | 1.3x | 1.3x | 1.2x | 1.2x | 1.2x | 1.2x |
| **DR** (Neo) | 1.1x | 1.0x | 1.0x | 1.0x | 1.1x | 1.1x | 1.1x | 1.0x | 1.1x | 1.1x | 1.0x | 1.0x | 1.1x | 1.1x | 1.1x | 1.1x |
| **DR** (both) | 2.4x | 2.6x | 2.7x | 3.2x | 2.5x | 2.7x | 2.9x | 2.5x | 3.4x | 3.2x | 3.1x | 3.5x | 2.8x | 2.4x | 2.5x | 3.0x |
| **NAND score** | 2.03 | 2.19 | 2.26 | 2.38 | 2.08 | 2.10 | 2.25 | 2.11 | 2.60 | 2.52 | 2.43 | 2.68 | 2.31 | 2.05 | 2.15 | 2.48 |

**b**

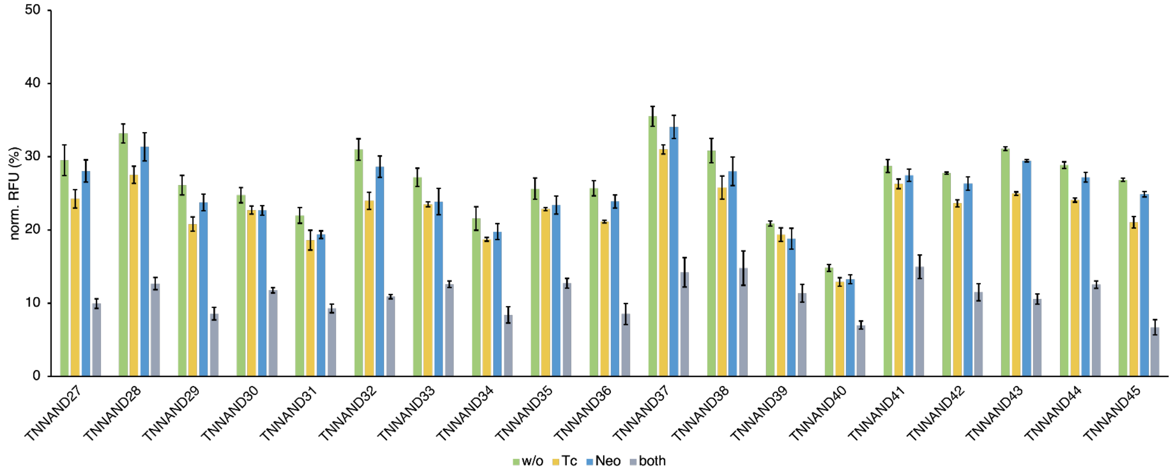

| | | | | | | | | | | | | | | | | | | |
|---|---|---|---|---|---|---|---|---|---|---|---|---|---|---|---|---|---|---|---|
| **DR** (Tc) | 1.2x | 1.2x | 1.3x | 1.1x | 1.2x | 1.3x | 1.2x | 1.2x | 1.1x | 1.2x | 1.1x | 1.2x | 1.1x | 1.1x | 1.1x | 1.2x | 1.2x | 1.2x | 1.3x |
| **DR** (Neo) | 1.1x | 1.1x | 1.1x | 1.1x | 1.1x | 1.1x | 1.1x | 1.1x | 1.1x | 1.1x | 1.0x | 1.1x | 1.1x | 1.1x | 1.0x | 1.1x | 1.1x | 1.1x | 1.1x |
| **DR** (both) | 3.0x | 2.6x | 3.0x | 2.1x | 2.4x | 2.8x | 2.2x | 2.6x | 2.0x | 3.0x | 2.5x | 2.1x | 1.8x | 2.1x | 1.9x | 2.4x | 2.9x | 2.3x | 4.0x |
| **NAND score** | 2.44 | 2.17 | 2.43 | 1.93 | 2.00 | 2.20 | 1.87 | 2.23 | 1.80 | 2.48 | 2.18 | 1.74 | 1.66 | 1.84 | 1.76 | 2.06 | 2.36 | 1.92 | 3.15 |

*Figure S3:* ***NAND machine learning training datasets.*** *Two sets of candidates (**a**) and (**b**) were chosen from the enriched TNNAND8 P1 sequence pool to be measured individually and serve as the initial training data for the ML algorithm. The influence on GFP expression of each P1 stem variant was measured in the absence (w/o) and presence of 250 µM tetracycline (Tc), 250 µM neomycin (Neo), or 250 µM of both. S. cerevisiae RS453 were transformed with the plasmid pcBB06 containing the respective candidate in the 5' UTR of a constitutively expressed gfp gene. Cells were incubated under the respective ligand conditions for 24 h. GFP values were normalized to constitutively expressed mCherry to eliminate cell to cell expression variability. pCBB06 without an insert was used as the blank and its values were subtracted from all other measurements. The fluorescence values of each measurement were normalized to its respective ligand-specific positive control (pcBB05). All measurements were performed in duplicates and repeated twice on separate days.*

### Figure S4 - ML Rounds 1-3 Datasets

**a**

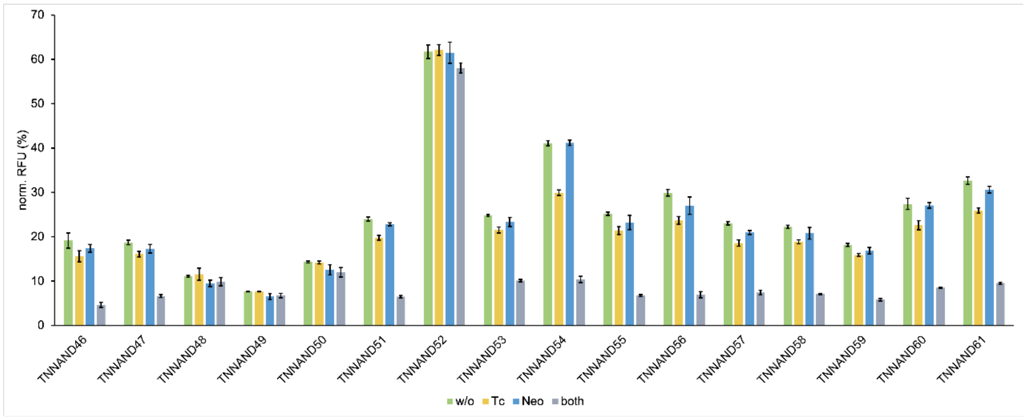

| | | | | | | | | | | | | | | | | |
|---|---|---|---|---|---|---|---|---|---|---|---|---|---|---|---|---|
| **DR** (Tc) | 1.2x | 1.2x | 1.0x | 1.0x | 1.0x | 1.2x | 1.0x | 1.2x | 1.4x | 1.2x | 1.3x | 1.2x | 1.2x | 1.1x | 1.2x | 1.3x |
| **DR** (Neo) | 1.1x | 1.1x | 1.2x | 1.2x | 1.1x | 1.1x | 1.0x | 1.1x | 1.0x | 1.1x | 1.1x | 1.1x | 1.1x | 1.1x | 1.0x | 1.1x |
| **DR** (both) | 4.1x | 2.8x | 1.1x | 1.1x | 1.2x | 3.7x | 1.1x | 2.5x | 4.0x | 3.7x | 4.3x | 3.1x | 3.1x | 3.1x | 3.2x | 3.4x |
| **NAND score** | 3.35 | 2.43 | 0.96 | 0.97 | 1.05 | 3.05 | 1.06 | 2.13 | 2.89 | 3.17 | 3.42 | 2.51 | 2.66 | 2.74 | 2.66 | 2.71 |

**b**

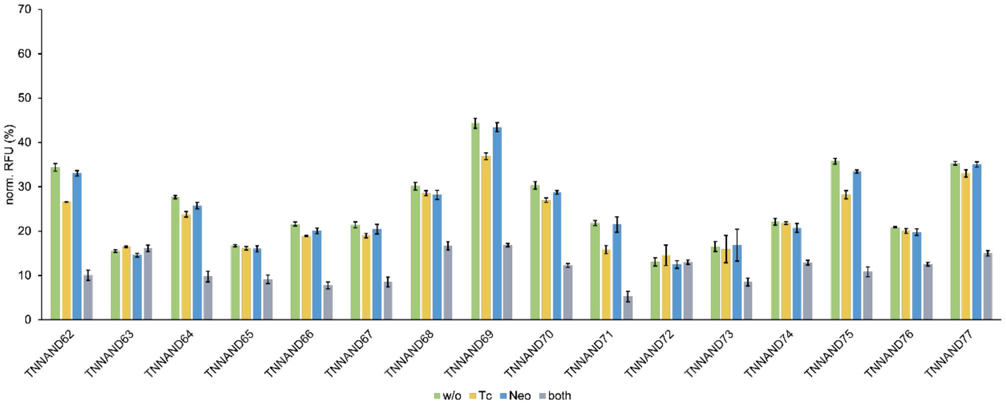

| | | | | | | | | | | | | | | | | |
|---|---|---|---|---|---|---|---|---|---|---|---|---|---|---|---|---|
| **DR** (Tc) | 1.3x | 0.9x | 1.2x | 1.0x | 1.1x | 1.1x | 1.1x | 1.2x | 1.1x | 1.4x | 0.9x | 1.0x | 1.0x | 1.3x | 1.0x | 1.1x |
| **DR** (Neo) | 1.0x | 1.1x | 1.1x | 1.0x | 1.1x | 1.0x | 1.1x | 1.0x | 1.1x | 1.0x | 1.0x | 1.0x | 1.1x | 1.1x | 1.1x | 1.0x |
| **DR** (both) | 3.4x | 1.0x | 2.8x | 1.8x | 2.8x | 2.5x | 1.8x | 2.6x | 2.5x | 4.1x | 1.0x | 1.9x | 1.7x | 3.3x | 1.7x | 2.4x |
| **NAND score** | 2.64 | 0.91 | 2.43 | 1.76 | 2.44 | 2.23 | 1.69 | 2.19 | 2.20 | 3.01 | 0.96 | 1.87 | 1.60 | 2.60 | 1.58 | 2.20 |

**c**

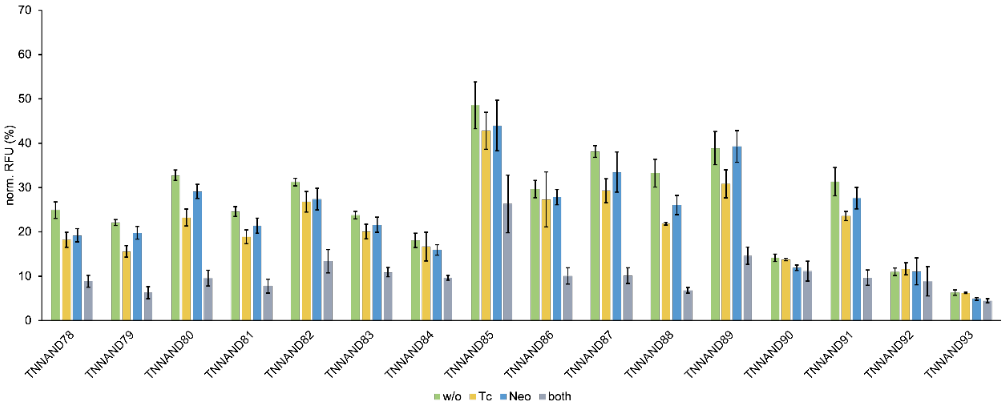

| | | | | | | | | | | | | | | | | |
|---|---|---|---|---|---|---|---|---|---|---|---|---|---|---|---|---|
| **DR** (Tc) | 1.4x | 1.4x | 1.4x | 1.3x | 1.2x | 1.2x | 1.1x | 1.1x | 1.1x | 1.3x | 1.5x | 1.3x | 1.0x | 1.3x | 0.9x | 1.0x |
| **DR** (Neo) | 1.3x | 1.1x | 1.1x | 1.1x | 1.1x | 1.1x | 1.1x | 1.1x | 1.1x | 1.1x | 1.3x | 1.0x | 1.2x | 1.1x | 1.0x | 1.3x |
| **DR** (both) | 2.8x | 3.5x | 3.4x | 3.2x | 2.3x | 2.2x | 1.9x | 1.8x | 2.9x | 3.8x | 4.9x | 2.7x | 1.3x | 3.2x | 1.2x | 1.4x |
| **NAND score** | 2.05 | 2.47 | 2.43 | 2.43 | 2.00 | 1.84 | 1.65 | 1.62 | 2.71 | 2.89 | 3.20 | 2.11 | 1.07 | 2.44 | 1.24 | 1.09 |

*Figure S4:* ***NAND machine learning rounds.*** *Three rounds of machine learning (**a**, **b** and **c**) were performed in total. Each batch of proposed candidates was measured individually to verify their switching behavior. The influence on GFP expression of each candidate was measured in the absence (w/o) and presence of 250 µM tetracycline (Tc), 250 µM neomycin (Neo), or 250 µM of both. S. cerevisiae RS453 were transformed with the plasmid pcBB06 containing the respective candidate in the 5' UTR of a constitutively expressed gfp gene. Cells were incubated under the respective ligand conditions for 24 h. GFP values were normalized to constitutively expressed mCherry to eliminate cell to cell expression variability. pCBB06 without an insert was used as the blank and its values were subtracted from all other measurements. The fluorescence values of each measurement were normalized to its respective ligand-specific positive control (pcBB05). All measurements were performed in duplicates and repeated twice on separate days.*

#### Table S1 - Construct Oligonucleotides for Hybridization

| ssDNA Oligonucleotide | Sequence (5' → 3') |
|---|---|
| TNNAND1_fw | CCGGTGCCTAAAACATACTAGCTTGTCCTTTAATGGTCCTAG AGAGGTGAAGAATACGACCACCTAGGCAAAATGG |
| TNNAND1_rev | CTAGCCATTTTGCCTAGGTGGTCGTATTCTTCACCTCTCTAG GACCATTAAAGGACAAGCTAGTATGTTTTAGGCA |
| TNNAND2_fw | CCGGTGCCTAAAACATACTGGCTTGTCCTTTAATGGTCCCG GAGAGGTGAAGAATACGACCACCTAGGCAAAATGG |
| TNNAND2_rev | CTAGCCATTTTGCCTAGGTGGTCGTATTCTTCACCTCTCCG GGACCATTAAAGGACAAGCCAGTATGTTTTAGGCA |
| TNNAND3_fw | CCGGTGCCTAAAACATACCGGCTTGTCCTTTAATGGTCCTG GAGAGGTGAAGAATACGACCACCTAGGCAAAATGG |
| TNNAND3_rev | CTAGCCATTTTGCCTAGGTGGTCGTATTCTTCACCTCTCCAG GACCATTAAAGGACAAGCCGGTATGTTTTAGGCA |
| TNNAND4_fw | CCGGTGCCTAAAACATACCCGCTTGTCCTTTAATGGTCCTG GAGAGGTGAAGAATACGACCACCTAGGCAAAATGG |
| TNNAND4_rev | CTAGCCATTTTGCCTAGGTGGTCGTATTCTTCACCTCTCCAG GACCATTAAAGGACAAGCGGGTATGTTTTAGGCA |
| TNNAND5_fw | CCGGTGCCTAAAACATACCTGCTTGTCCTTTAATGGTCCCG GAGAGGTGAAGAATACGACCACCTAGGCAAAATGG |
| TNNAND5_rev | CTAGCCATTTTGCCTAGGTGGTCGTATTCTTCACCTCTCCG GGACCATTAAAGGACAAGCAGGTATGTTTTAGGCA |
| TNNAND6_fw | CCGGTGCCTAAAACATACCCGCTTGTCCTTTAATGGTCCGT GAGAGGTGAAGAATACGACCACCTAGGCAAAATGG |
| TNNAND6_rev | CTAGCCATTTTGCCTAGGTGGTCGTATTCTTCACCTCTCACG GACCATTAAAGGACAAGCGGGTATGTTTTAGGCA |
| TNNAND7_fw | CCGGTGCCTAAAACATAGTCGCTTGTCCTTTAATGGTCCGT CAGAGGTGAAGAATACGACCACCTAGGCAAAATGG |

| | |
|---|---|
| TNNAND7_rev | CTAGCCATTTTGCCTAGGTGGTCGTATTCTTCACCTCTGAC GGACCATTAAAGGACAAGCGACTATGTTTTAGGCA |
| TNNAND8_fw | CCGGTGCCTAAAACATACTCGCTTGTCCTTTAATGGTCCTTG AGAGGTGAAGAATACGACCACCTAGGCAAAATGG |
| TNNAND8_rev | CTAGCCATTTTGCCTAGGTGGTCGTATTCTTCACCTCTCAAG GACCATTAAAGGACAAGCGAGTATGTTTTAGGCA |
| TNNAND9_fw | CCGGTGCCTAAAACATACCCGCTTGTCCTTTAATGGTCCGA GAGAGGTGAAGAATACGACCACCTAGGCAAAATGG |
| TNNAND9_rev | CTAGCCATTTTGCCTAGGTGGTCGTATTCTTCACCTCTCTCG GACCATTAAAGGACAAGCGGGTATGTTTTAGGCA |
| TNNAND10_fw | CCGGTGCCTAAAACATACTAGCTTGTCCTTTAATGGTCCTG GAGAGGTGAAGAATACGACCACCTAGGCAAAATGG |
| TNNAND10_rev | CTAGCCATTTTGCCTAGGTGGTCGTATTCTTCACCTCTCCAG GACCATTAAAGGACAAGCTAGTATGTTTTAGGCA |
| TNNAND11_fw | CCGGTTGCCCGCAAACATACTCGCTTGTCCTTTAATGGTCC TTGAGAGGTGAAGAATACGACCACCGCGGGTTAAAATGG |
| TNNAND11_rev | CTAGCCATTTTAACCCGCGGTGGTCGTATTCTTCACCTCTCA AGGACCATTAAAGGACAAGCGAGTATGTTTGCGGGCAA |
| TNNAND12_fw | CCGGTAGACGCCAAACATACTCGCTTGTCCTTTAATGGTCC TTGAGAGGTGAAGAATACGACCACCGGCGTTGAAAATGG |
| TNNAND12_rev | CTAGCCATTTTCAACGCCGGTGGTCGTATTCTTCACCTCTCA AGGACCATTAAAGGACAAGCGAGTATGTTTGGCGTCTA |
| TNNAND13_fw | CCGGTAGCGCGCAAACATACTCGCTTGTCCTTTAATGGTCC TTGAGAGGTGAAGAATACGACCACCGCGTGCGAAAATGG |
| TNNAND13_rev | CTAGCCATTTTCGCACGCGGTGGTCGTATTCTTCACCTCTC AAGGACCATTAAAGGACAAGCGAGTATGTTTGCGCGCTA |
| TNNAND14_fw | CCGGTTACCCGCAAACATACTCGCTTGTCCTTTAATGGTCC TTGAGAGGTGAAGAATACGACCACCGCGGAGTAAAATGG |
| TNNAND14_rev | CTAGCCATTTTACTCCGCGGTGGTCGTATTCTTCACCTCTCA AGGACCATTAAAGGACAAGCGAGTATGTTTGCGGGTAA |

| | |
|---|---|
| TNNAND15_fw | CCGGTACCCTGCAAACATACTCGCTTGTCCTTTAATGGTCC TTGAGAGGTGAAGAATACGACCACCGCAGGGAAAAATGG |
| TNNAND15_rev | CTAGCCATTTTTCCCTGCGGTGGTCGTATTCTTCACCTCTCA AGGACCATTAAAGGACAAGCGAGTATGTTTGCAGGGTA |
| TNNAND16_fw | CCGGTTTCGCGCAAACATACTCGCTTGTCCTTTAATGGTCC TTGAGAGGTGAAGAATACGACCACCGCGCGGGAAAATGG |
| TNNAND16_rev | CTAGCCATTTTCCCGCGCGGTGGTCGTATTCTTCACCTCTC AAGGACCATTAAAGGACAAGCGAGTATGTTTGCGCGAAA |
| TNNAND17_fw | CCGGTTGCGCGCAAACATACTCGCTTGTCCTTTAATGGTCC TTGAGAGGTGAAGAATACGACCACCGCGCGTAAAAATGG |
| TNNAND17_rev | CTAGCCATTTTTACGCGCGGTGGTCGTATTCTTCACCTCTCA AGGACCATTAAAGGACAAGCGAGTATGTTTGCGCGCAA |
| TNNAND18_fw | CCGGTCGCGTGCAAACATACTCGCTTGTCCTTTAATGGTCC TTGAGAGGTGAAGAATACGACCACCGCACGTGAAAATGG |
| TNNAND18_rev | CTAGCCATTTTCACGTGCGGTGGTCGTATTCTTCACCTCTCA AGGACCATTAAAGGACAAGCGAGTATGTTTGCACGCGA |
| TNNAND19_fw | CCGGTAGCGCGCAAACATACTCGCTTGTCCTTTAATGGTCC TTGAGAGGTGAAGAATACGACCACCGCGCGCGAAAATGG |
| TNNAND19_rev | CTAGCCATTTTCGCGCGCGGTGGTCGTATTCTTCACCTCTC AAGGACCATTAAAGGACAAGCGAGTATGTTTGCGCGCTA |
| TNNAND20_fw | CCGGTTCCCAGCAAACATACTCGCTTGTCCTTTAATGGTCC TTGAGAGGTGAAGAATACGACCACCGCTGGGTAAAATGG |
| TNNAND20_rev | CTAGCCATTTTACCCAGCGGTGGTCGTATTCTTCACCTCTCA AGGACCATTAAAGGACAAGCGAGTATGTTTGCTGGGAA |
| TNNAND21_fw | CCGGTTTCCCGCAAACATACTCGCTTGTCCTTTAATGGTCCT TGAGAGGTGAAGAATACGACCACCGCGGGGGAAAATGG |
| TNNAND21_rev | CTAGCCATTTTCCCCCGCGGTGGTCGTATTCTTCACCTCTC AAGGACCATTAAAGGACAAGCGAGTATGTTTGCGGGAAA |
| TNNAND22_fw | CCGGTCTGCGCCAAACATACTCGCTTGTCCTTTAATGGTCC TTGAGAGGTGAAGAATACGACCACCGGCGCGGAAAATGG |

| | |
|---|---|
| TNNAND22_rev | CTAGCCATTTTCCGCGCCGGTGGTCGTATTCTTCACCTCTCAAGGACCATTAAAGGACAAGCGAGTATGTTTGGCGCAGA |
| TNNAND23_fw | CCGGTAACCCGCAAACATACTCGCTTGTCCTTTAATGGTCCTTGAGAGGTGAAGAATACGACCACCGCGGGTAAAAATGG |
| TNNAND23_rev | CTAGCCATTTTTACCCGCGGTGGTCGTATTCTTCACCTCTCAAGGACCATTAAAGGACAAGCGAGTATGTTTGCGGGTTA |
| TNNAND24_fw | CCGGTCGCAGCCAAACATACTCGCTTGTCCTTTAATGGTCCTTGAGAGGTGAAGAATACGACCACCGGCTGTGAAAATGG |
| TNNAND24_rev | CTAGCCATTTTCACAGCCGGTGGTCGTATTCTTCACCTCTCAAGGACCATTAAAGGACAAGCGAGTATGTTTGGCTGCGA |
| TNNAND25_fw | CCGGTGTCACGCAAACATACTCGCTTGTCCTTTAATGGTCCTTGAGAGGTGAAGAATACGACCACCGCGTGACAAAATGG |
| TNNAND25_rev | CTAGCCATTTTGTCACGCGGTGGTCGTATTCTTCACCTCTCAAGGACCATTAAAGGACAAGCGAGTATGTTTGCGTGACA |
| TNNAND26_fw | CCGGTGCTCCGCAAACATACTCGCTTGTCCTTTAATGGTCCTTGAGAGGTGAAGAATACGACCACCGCGGAGTAAAATGG |
| TNNAND26_rev | CTAGCCATTTTACTCCGCGGTGGTCGTATTCTTCACCTCTCAAGGACCATTAAAGGACAAGCGAGTATGTTTGCGGAGCA |
| TNNAND27_fw | CCGGTCGCGTGCAAACATACTCGCTTGTCCTTTAATGGTCCTTGAGAGGTGAAGAATACGACCACCGCACGCGAAAATGG |
| TNNAND27_rev | CTAGCCATTTTCGCGTGCGGTGGTCGTATTCTTCACCTCTCAAGGACCATTAAAGGACAAGCGAGTATGTTTGCACGCGA |
| TNNAND28_fw | CCGGTTACCCGCAAACATACTCGCTTGTCCTTTAATGGTCCTTGAGAGGTGAAGAATACGACCACCGCGGGGTAAAATGG |
| TNNAND28_rev | CTAGCCATTTTACCCCGCGGTGGTCGTATTCTTCACCTCTCAAGGACCATTAAAGGACAAGCGAGTATGTTTGCGGGTAA |
| TNNAND29_fw | CCGGTCCTCTGCAAACATACTCGCTTGTCCTTTAATGGTCCTTGAGAGGTGAAGAATACGACCACCGCAGGGGAAAATGG |
| TNNAND29_rev | CTAGCCATTTTCCCCTGCGGTGGTCGTATTCTTCACCTCTCAAGGACCATTAAAGGACAAGCGAGTATGTTTGCAGAGGA |

| | |
|---|---|
| TNNAND30_fw | CCGGTTTCGCCCAAACATACTCGCTTGTCCTTTAATGGTCCTTGAGAGGTGAAGAATACGACCACCGGGCGGGAAAATGG |
| TNNAND30_rev | CTAGCCATTTTCCCGCCCGGTGGTCGTATTCTTCACCTCTCAAGGACCATTAAAGGACAAGCGAGTATGTTTGGGCGAAA |
| TNNAND31_fw | CCGGTTACCGCCAAACATACTCGCTTGTCCTTTAATGGTCCTTGAGAGGTGAAGAATACGACCACCGGCGGCAAAAATGG |
| TNNAND31_rev | CTAGCCATTTTTGCCGCCGGTGGTCGTATTCTTCACCTCTCAAGGACCATTAAAGGACAAGCGAGTATGTTTGGCGGTAA |
| TNNAND32_fw | CCGGTTTCCGCCAAACATACTCGCTTGTCCTTTAATGGTCCTTGAGAGGTGAAGAATACGACCACCGGCGCGAAAAATGG |
| TNNAND32_rev | CTAGCCATTTTTCGCGCCGGTGGTCGTATTCTTCACCTCTCAAGGACCATTAAAGGACAAGCGAGTATGTTTGGCGGAAA |
| TNNAND33_fw | CCGGTAGCCAGCAAACATACTCGCTTGTCCTTTAATGGTCCTTGAGAGGTGAAGAATACGACCACCGCTGGGTAAAATGG |
| TNNAND33_rev | CTAGCCATTTTACCCAGCGGTGGTCGTATTCTTCACCTCTCAAGGACCATTAAAGGACAAGCGAGTATGTTTGCTGGCTA |
| TNNAND34_fw | CCGGTTGCACCCAAACATACTCGCTTGTCCTTTAATGGTCCTTGAGAGGTGAAGAATACGACCACCGGGTGCTAAAATGG |
| TNNAND34_rev | CTAGCCATTTTAGCACCCGGTGGTCGTATTCTTCACCTCTCAAGGACCATTAAAGGACAAGCGAGTATGTTTGGGTGCAA |
| TNNAND35_fw | CCGGTCCTGCGCAAACATACTCGCTTGTCCTTTAATGGTCCTTGAGAGGTGAAGAATACGACCACCGCCAGGCAAAATGG |
| TNNAND35_rev | CTAGCCATTTTGCCTGGCGGTGGTCGTATTCTTCACCTCTCAAGGACCATTAAAGGACAAGCGAGTATGTTTGCGCAGGA |
| TNNAND36_fw | CCGGTACGCGGCAAACATACTCGCTTGTCCTTTAATGGTCCTTGAGAGGTGAAGAATACGACCACCGCCGCGCAAAATGG |
| TNNAND36_rev | CTAGCCATTTTGCGCGGCGGTGGTCGTATTCTTCACCTCTCAAGGACCATTAAAGGACAAGCGAGTATGTTTGCCGCGTA |
| TNNAND37_fw | CCGGTCTCCCGCAAACATACTCGCTTGTCCTTTAATGGTCCTTGAGAGGTGAAGAATACGACCACCGCGGGGTAAAATGG |

| | |
|---|---|
| TNNAND37_rev | CTAGCCATTTTACCCCGCGGTGGTCGTATTCTTCACCTCTC AAGGACCATTAAAGGACAAGCGAGTATGTTTGCGGGAGA |
| TNNAND38_fw | CCGGTTTTCGGCAAACATACTCGCTTGTCCTTTAATGGTCCT TGAGAGGTGAAGAATACGACCACCGCCGTCCAAAATGG |
| TNNAND38_rev | CTAGCCATTTTGGACGGCGGTGGTCGTATTCTTCACCTCTC AAGGACCATTAAAGGACAAGCGAGTATGTTTGCCGAAAA |
| TNNAND39_fw | CCGGTTGCCCCAAACATACTCGCTTGTCCTTTAATGGTCCTT GAGAGGTGAAGAATACGACCACCGGGGCTAAAATGG |
| TNNAND39_rev | CTAGCCATTTTAGCCCCGGTGGTCGTATTCTTCACCTCTCAA GGACCATTAAAGGACAAGCGAGTATGTTTGGGGCAA |
| TNNAND40_fw | CCGGTTCTCCCCAAACATACTCGCTTGTCCTTTAATGGTCCT TGAGAGGTGAAGAATACGACCACCGGGGTGGAAAATGG |
| TNNAND40_rev | CTAGCCATTTTCCACCCCGGTGGTCGTATTCTTCACCTCTCA AGGACCATTAAAGGACAAGCGAGTATGTTTGGGGAGAA |
| TNNAND41_fw | CCGGTTCGGGGCAAACATACTCGCTTGTCCTTTAATGGTCC TTGAGAGGTGAAGAATACGACCACCGCCCTGGAAAATGG |
| TNNAND41_rev | CTAGCCATTTTCCAGGGCGGTGGTCGTATTCTTCACCTCTC AAGGACCATTAAAGGACAAGCGAGTATGTTTGCCCCGAA |
| TNNAND42_fw | CCGGTCCCTGCCAAACATACTCGCTTGTCCTTTAATGGTCC TTGAGAGGTGAAGAATACGACCACCGGGGGGGAAAATGG |
| TNNAND42_rev | CTAGCCATTTTCCCCCCCGGTGGTCGTATTCTTCACCTCTC AAGGACCATTAAAGGACAAGCGAGTATGTTTGGCAGGGA |
| TNNAND43_fw | CCGGTCTCGCGCAAACATACTCGCTTGTCCTTTAATGGTCC TTGAGAGGTGAAGAATACGACCACCGCGCGGGAAAATGG |
| TNNAND43_rev | CTAGCCATTTTCCCGCGCGGTGGTCGTATTCTTCACCTCTC AAGGACCATTAAAGGACAAGCGAGTATGTTTGCGCGAGA |
| TNNAND44_fw | CCGGTCACGTCAAACATACTCGCTTGTCCTTTAATGGTCCTT GAGAGGTGAAGAATACGACCACCGGCGTGAAAATGG |
| TNNAND44_rev | CTAGCCATTTTCACGCCGGTGGTCGTATTCTTCACCTCTCAA GGACCATTAAAGGACAAGCGAGTATGTTTGACGTGA |

| | |
|---|---|
| TNNAND45_fw | CCGGTCCTCTGCAAACATACTCGCTTGTCCTTTAATGGTCCTTGAGAGGTGAAGAATACGACCACCGCAGGGGAAAATGG |
| TNNAND45_rev | CTAGCCATTTTCCCCTGCGGTGGTCGTATTCTTCACCTCTCAAGGACCATTAAAGGACAAGCGAGTATGTTTGCAGAGGA |
| TNNAND46_fw | CCGGTGCCTGCAAACATACTCGCTTGTCCTTTAATGGTCCTTGAGAGGTGAAGAATACGACCACCGCAGGCAAAATGG |
| TNNAND46_rev | CTAGCCATTTTGCCTGCGGTGGTCGTATTCTTCACCTCTCAAGGACCATTAAAGGACAAGCGAGTATGTTTGCAGGCA |
| TNNAND47_fw | CCGGTTCGCTGCAAACATACTCGCTTGTCCTTTAATGGTCCTTGAGAGGTGAAGAATACGACCACCGCAGTGGAAAATGG |
| TNNAND47_rev | CTAGCCATTTTCCACTGCGGTGGTCGTATTCTTCACCTCTCAAGGACCATTAAAGGACAAGCGAGTATGTTTGCAGCGAA |
| TNNAND48_fw | CCGGTGCGATAAACATACTCGCTTGTCCTTTAATGGTCCTTGAGAGGTGAAGAATACGACCACCATGCAAAAATGG |
| TNNAND48_rev | CTAGCCATTTTTGCATGGTGGTCGTATTCTTCACCTCTCAAGGACCATTAAAGGACAAGCGAGTATGTTTATCGCA |
| TNNAND49_fw | CCGGTCGGAACAAACATACTCGCTTGTCCTTTAATGGTCCTTGAGAGGTGAAGAATACGACCACCGTCGTAAAAATGG |
| TNNAND49_rev | CTAGCCATTTTTACGACGGTGGTCGTATTCTTCACCTCTCAAGGACCATTAAAGGACAAGCGAGTATGTTTGTTCCGA |
| TNNAND50_fw | CCGGTTAATGAAACATACTCGCTTGTCCTTTAATGGTCCTTGAGAGGTGAAGAATACGACCACCAAGACAAAATGG |
| TNNAND50_rev | CTAGCCATTTTGTCTTGGTGGTCGTATTCTTCACCTCTCAAGGACCATTAAAGGACAAGCGAGTATGTTTCATTAA |
| TNNAND51_fw | CCGGTCTGTCCCAAACATACTCGCTTGTCCTTTAATGGTCCTTGAGAGGTGAAGAATACGACCACCGGGACGGAAAATGG |
| TNNAND51_rev | CTAGCCATTTTCCGTCCCGGTGGTCGTATTCTTCACCTCTCAAGGACCATTAAAGGACAAGCGAGTATGTTTGGGACAGA |
| TNNAND52_fw | CCGGTCTTGTAAACATACTCGCTTGTCCTTTAATGGTCCTTGAGAGGTGAAGAATACGACCACCCTGGAAAAATGG |

| | |
|---|---|
| TNNAND52_rev | CTAGCCATTTTTCCAGGGTGGTCGTATTCTTCACCTCTCAAG GACCATTAAAGGACAAGCGAGTATGTTTACAAGA |
| TNNAND53_fw | CCGGTCCCCTGCAAACATACTCGCTTGTCCTTTAATGGTCC TTGAGAGGTGAAGAATACGACCACCGCTAGGGAAAATGG |
| TNNAND53_rev | CTAGCCATTTTCCCTAGCGGTGGTCGTATTCTTCACCTCTCA AGGACCATTAAAGGACAAGCGAGTATGTTTGCAGGGGA |
| TNNAND54_fw | CCGGTCCCGTGCAAACATACTCGCTTGTCCTTTAATGGTCC TTGAGAGGTGAAGAATACGACCACCGCAGGGGAAAATGG |
| TNNAND54_rev | CTAGCCATTTTCCCCTGCGGTGGTCGTATTCTTCACCTCTCA AGGACCATTAAAGGACAAGCGAGTATGTTTGCACGGGA |
| TNNAND55_fw | CCGGTAGGGAGCAAACATACTCGCTTGTCCTTTAATGGTCC TTGAGAGGTGAAGAATACGACCACCGCTCCCAAAAATGG |
| TNNAND55_rev | CTAGCCATTTTTGGGAGCGGTGGTCGTATTCTTCACCTCTC AAGGACCATTAAAGGACAAGCGAGTATGTTTGCTCCCTA |
| TNNAND56_fw | CCGGTAACTCCCAAACATACTCGCTTGTCCTTTAATGGTCCT TGAGAGGTGAAGAATACGACCACCGGGAGCGAAAATGG |
| TNNAND56_rev | CTAGCCATTTTCGCTCCCGGTGGTCGTATTCTTCACCTCTCA AGGACCATTAAAGGACAAGCGAGTATGTTTGGGAGTTA |
| TNNAND57_fw | CCGGTCCACCCCAAACATACTCGCTTGTCCTTTAATGGTCC TTGAGAGGTGAAGAATACGACCACCGGGTTGGAAAATGG |
| TNNAND57_rev | CTAGCCATTTTCCAACCCGGTGGTCGTATTCTTCACCTCTCA AGGACCATTAAAGGACAAGCGAGTATGTTTGGGGTGGA |
| TNNAND58_fw | CCGGTACGTTGCAAACATACTCGCTTGTCCTTTAATGGTCCT TGAGAGGTGAAGAATACGACCACCGCAGCGGAAAATGG |
| TNNAND58_rev | CTAGCCATTTTCCGCTGCGGTGGTCGTATTCTTCACCTCTC AAGGACCATTAAAGGACAAGCGAGTATGTTTGCAACGTA |
| TNNAND59_fw | CCGGTGTGCGCCAAACATACTCGCTTGTCCTTTAATGGTCC TTGAGAGGTGAAGAATACGACCACCGGCGCGCAAAATGG |
| TNNAND59_rev | CTAGCCATTTTGCGCGCCGGTGGTCGTATTCTTCACCTCTC AAGGACCATTAAAGGACAAGCGAGTATGTTTGGCGCACA |

| | |
|---|---|
| TNNAND60_fw | CCGGTTTGCTGCAAACATACTCGCTTGTCCTTTAATGGTCCTTGAGAGGTGAAGAATACGACCACCGCAGCATAAAATGG |
| TNNAND60_rev | CTAGCCATTTTATGCTGCGGTGGTCGTATTCTTCACCTCTCAAGGACCATTAAAGGACAAGCGAGTATGTTTGCAGCAAA |
| TNNAND61_fw | CCGGTTATCAGCAAACATACTCGCTTGTCCTTTAATGGTCCTTGAGAGGTGAAGAATACGACCACCGCTGGTAAAAATGG |
| TNNAND61_rev | CTAGCCATTTTTACCAGCGGTGGTCGTATTCTTCACCTCTCAAGGACCATTAAAGGACAAGCGAGTATGTTTGCTGATAA |
| TNNAND62_fw | CCGGTTCCTGCCAAACATACTCGCTTGTCCTTTAATGGTCCTTGAGAGGTGAAGAATACGACCACCGGCAAGGAAAATGG |
| TNNAND62_rev | CTAGCCATTTTCCTTGCCGGTGGTCGTATTCTTCACCTCTCAAGGACCATTAAAGGACAAGCGAGTATGTTTGGCAGGAA |
| TNNAND63_fw | CCGGTGTAGACCAAACATACTCGCTTGTCCTTTAATGGTCCTTGAGAGGTGAAGAATACGACCACCGGAATGGAAAATGG |
| TNNAND63_rev | CTAGCCATTTTCCATTCCGGTGGTCGTATTCTTCACCTCTCAAGGACCATTAAAGGACAAGCGAGTATGTTTGGTCTACA |
| TNNAND64_fw | CCGGTGGGGTGCAAACATACTCGCTTGTCCTTTAATGGTCCTTGAGAGGTGAAGAATACGACCACCGCATCCCAAAATGG |
| TNNAND64_rev | CTAGCCATTTTGGGATGCGGTGGTCGTATTCTTCACCTCTCAAGGACCATTAAAGGACAAGCGAGTATGTTTGCACCCCA |
| TNNAND65_fw | CCGGTGCGGACCAAACATACTCGCTTGTCCTTTAATGGTCCTTGAGAGGTGAAGAATACGACCACCGGCCCGCAAAATGG |
| TNNAND65_rev | CTAGCCATTTTGCGGGCCGGTGGTCGTATTCTTCACCTCTCAAGGACCATTAAAGGACAAGCGAGTATGTTTGGTCCGCA |
| TNNAND66_fw | CCGGTGCCTTCCAAACATACTCGCTTGTCCTTTAATGGTCCTTGAGAGGTGAAGAATACGACCACCGGAAGGCAAAATGG |
| TNNAND66_rev | CTAGCCATTTTGCCTTCCGGTGGTCGTATTCTTCACCTCTCAAGGACCATTAAAGGACAAGCGAGTATGTTTGGAAGGCA |
| TNNAND67_fw | CCGGTAAGTCGCAAACATACTCGCTTGTCCTTTAATGGTCCTTGAGAGGTGAAGAATACGACCACCGCGACCGAAAATGG |

| | |
|---|---|
| TNNAND67_rev | CTAGCCATTTTCGGTCGCGGTGGTCGTATTCTTCACCTCTCAAGGACCATTAAAGGACAAGCGAGTATGTTTGCGACTTA |
| TNNAND68_fw | CCGGTGCTTGCAAACATACTCGCTTGTCCTTTAATGGTCCTTGAGAGGTGAAGAATACGACCACCGCAAGCAAAATGG |
| TNNAND68_rev | CTAGCCATTTTGCTTGCGGTGGTCGTATTCTTCACCTCTCAAGGACCATTAAAGGACAAGCGAGTATGTTTGCAAGCA |
| TNNAND69_fw | CCGGTCTGCCCCAAACATACTCGCTTGTCCTTTAATGGTCCTTGAGAGGTGAAGAATACGACCACCGGAGCAGAAAATGG |
| TNNAND69_rev | CTAGCCATTTTCTGCTCCGGTGGTCGTATTCTTCACCTCTCAAGGACCATTAAAGGACAAGCGAGTATGTTTGGGGCAGA |
| TNNAND70_fw | CCGGTTAGTCCCAAACATACTCGCTTGTCCTTTAATGGTCCTTGAGAGGTGAAGAATACGACCACCGGGACTAAAAATGG |
| TNNAND70_rev | CTAGCCATTTTTAGTCCCGGTGGTCGTATTCTTCACCTCTCAAGGACCATTAAAGGACAAGCGAGTATGTTTGGGACTAA |
| TNNAND71_fw | CCGGTAGCACCCAAACATACTCGCTTGTCCTTTAATGGTCCTTGAGAGGTGAAGAATACGACCACCGGGTGCGAAAATGG |
| TNNAND71_rev | CTAGCCATTTTCGCACCCGGTGGTCGTATTCTTCACCTCTCAAGGACCATTAAAGGACAAGCGAGTATGTTTGGGTGCTA |
| TNNAND72_fw | CCGGTGTGACCCAAACATACTCGCTTGTCCTTTAATGGTCCTTGAGAGGTGAAGAATACGACCACCGGAGCCCAAAATGG |
| TNNAND72_rev | CTAGCCATTTTGGGCTCCGGTGGTCGTATTCTTCACCTCTCAAGGACCATTAAAGGACAAGCGAGTATGTTTGGGTCACA |
| TNNAND73_fw | CCGGTGCCTTGCAAACATACTCGCTTGTCCTTTAATGGTCCTTGAGAGGTGAAGAATACGACCACCGCTTGGCAAAATGG |
| TNNAND73_rev | CTAGCCATTTTGCCAAGCGGTGGTCGTATTCTTCACCTCTCAAGGACCATTAAAGGACAAGCGAGTATGTTTGCAAGGCA |
| TNNAND74_fw | CCGGTCTGTCCCAAACATACTCGCTTGTCCTTTAATGGTCCTTGAGAGGTGAAGAATACGACCACCGGGCGGCAAAATGG |
| TNNAND74_rev | CTAGCCATTTTGCCGCCCGGTGGTCGTATTCTTCACCTCTCAAGGACCATTAAAGGACAAGCGAGTATGTTTGGGACAGA |

| | |
|---|---|
| TNNAND75_fw | CCGGTCCTTACCAAACATACTCGCTTGTCCTTTAATGGTCCT TGAGAGGTGAAGAATACGACCACCGGTAAGGAAAATGG |
| TNNAND75_rev | CTAGCCATTTTCCTTACCGGTGGTCGTATTCTTCACCTCTCA AGGACCATTAAAGGACAAGCGAGTATGTTTGGTAAGGA |
| TNNAND76_fw | CCGGTGCCTGCCAAACATACTCGCTTGTCCTTTAATGGTCC TTGAGAGGTGAAGAATACGACCACCGGCAGCAAAAATGG |
| TNNAND76_rev | CTAGCCATTTTTGCTGCCGGTGGTCGTATTCTTCACCTCTCA AGGACCATTAAAGGACAAGCGAGTATGTTTGGCAGGCA |
| TNNAND77_fw | CCGGTCGCTCCCAAACATACTCGCTTGTCCTTTAATGGTCC TTGAGAGGTGAAGAATACGACCACCGGGAGCAAAAATGG |
| TNNAND77_rev | CTAGCCATTTTTGCTCCCGGTGGTCGTATTCTTCACCTCTCA AGGACCATTAAAGGACAAGCGAGTATGTTTGGGAGCGA |
| TNNAND78_fw | CCGGTACTTCCCAAACATACTCGCTTGTCCTTTAATGGTCCT TGAGAGGTGAAGAATACGACCACCGGGGGGGAAAATGG |
| TNNAND78_rev | CTAGCCATTTTCCCCCCCGGTGGTCGTATTCTTCACCTCTC AAGGACCATTAAAGGACAAGCGAGTATGTTTGGGAAGTA |
| TNNAND79_fw | CCGGTTCCACCCAAACATACTCGCTTGTCCTTTAATGGTCCT TGAGAGGTGAAGAATACGACCACCGGGTGCGAAAATGG |
| TNNAND79_rev | CTAGCCATTTTCGCACCCGGTGGTCGTATTCTTCACCTCTC AAGGACCATTAAAGGACAAGCGAGTATGTTTGGGTGGAA |
| TNNAND80_fw | CCGGTTCCGTGCAAACATACTCGCTTGTCCTTTAATGGTCC TTGAGAGGTGAAGAATACGACCACCGCACAGGAAAATGG |
| TNNAND80_rev | CTAGCCATTTTCCTGTGCGGTGGTCGTATTCTTCACCTCTCA AGGACCATTAAAGGACAAGCGAGTATGTTTGCACGGAA |
| TNNAND81_fw | CCGGTCGGTTGCAAACATACTCGCTTGTCCTTTAATGGTCC TTGAGAGGTGAAGAATACGACCACCGCAACCGAAAATGG |
| TNNAND81_rev | CTAGCCATTTTCGGTTGCGGTGGTCGTATTCTTCACCTCTCA AGGACCATTAAAGGACAAGCGAGTATGTTTGCAACCGA |
| TNNAND82_fw | CCGGTCATCCGCAAACATACTCGCTTGTCCTTTAATGGTCC TTGAGAGGTGAAGAATACGACCACCGCGGGCAAAATGG |

| | |
|---|---|
| TNNAND82_rev | CTAGCCATTTTGCCCCGCGGTGGTCGTATTCTTCACCTCTC AAGGACCATTAAAGGACAAGCGAGTATGTTTGCGGATGA |
| TNNAND83_fw | CCGGTAGCGTCCAAACATACTCGCTTGTCCTTTAATGGTCC TTGAGAGGTGAAGAATACGACCACCGGGCGCGAAAATGG |
| TNNAND83_rev | CTAGCCATTTTCGCGCCCGGTGGTCGTATTCTTCACCTCTC AAGGACCATTAAAGGACAAGCGAGTATGTTTGGACGCTA |
| TNNAND84_fw | CCGGTGCCTGCAAACATACTCGCTTGTCCTTTAATGGTCCT TGAGAGGTGAAGAATACGACCACCGCGGGCAAAATGG |
| TNNAND84_rev | CTAGCCATTTTGCCCGCGGTGGTCGTATTCTTCACCTCTCA AGGACCATTAAAGGACAAGCGAGTATGTTTGCAGGCA |
| TNNAND85_fw | CCGGTTCCTACCAAACATACTCGCTTGTCCTTTAATGGTCCT TGAGAGGTGAAGAATACGACCACCGGTACGGAAAATGG |
| TNNAND85_rev | CTAGCCATTTTCCGTACCGGTGGTCGTATTCTTCACCTCTCA AGGACCATTAAAGGACAAGCGAGTATGTTTGGTAGGAA |
| TNNAND86_fw | CCGGTCGGGTGCAAACATACTCGCTTGTCCTTTAATGGTCC TTGAGAGGTGAAGAATACGACCACCGCACCCCAAAATGG |
| TNNAND86_rev | CTAGCCATTTTGGGGTGCGGTGGTCGTATTCTTCACCTCTC AAGGACCATTAAAGGACAAGCGAGTATGTTTGCACCCGA |
| TNNAND87_fw | CCGGTCTGCTGCAAACATACTCGCTTGTCCTTTAATGGTCC TTGAGAGGTGAAGAATACGACCACCGCAGCGGAAAATGG |
| TNNAND87_rev | CTAGCCATTTTCCGCTGCGGTGGTCGTATTCTTCACCTCTC AAGGACCATTAAAGGACAAGCGAGTATGTTTGCAGCAGA |
| TNNAND88_fw | CCGGTCTGCACCAAACATACTCGCTTGTCCTTTAATGGTCC TTGAGAGGTGAAGAATACGACCACCGGTGCAGAAAATGG |
| TNNAND88_rev | CTAGCCATTTTCTGCACCGGTGGTCGTATTCTTCACCTCTCA AGGACCATTAAAGGACAAGCGAGTATGTTTGGTGCAGA |
| TNNAND89_fw | CCGGTCTGTGCCAAACATACTCGCTTGTCCTTTAATGGTCC TTGAGAGGTGAAGAATACGACCACCGGCACCGAAAATGG |
| TNNAND89_rev | CTAGCCATTTTCGGTGCCGGTGGTCGTATTCTTCACCTCTC AAGGACCATTAAAGGACAAGCGAGTATGTTTGGCACAGA |

| | |
|---|---|
| TNNAND90_fw | CCGGTTGGCCCCAAACATACTCGCTTGTCCTTTAATGGTCC TTGAGAGGTGAAGAATACGACCACCGGGGACGAAAATGG |
| TNNAND90_rev | CTAGCCATTTTCGTCCCCGGTGGTCGTATTCTTCACCTCTCA AGGACCATTAAAGGACAAGCGAGTATGTTTGGGGCCAA |
| TNNAND91_fw | CCGGTTGGGAGCAAACATACTCGCTTGTCCTTTAATGGTCC TTGAGAGGTGAAGAATACGACCACCGCTCCCGAAAATGG |
| TNNAND91_rev | CTAGCCATTTTCGGGAGCGGTGGTCGTATTCTTCACCTCTC AAGGACCATTAAAGGACAAGCGAGTATGTTTGCTCCCAA |
| TNNAND92_fw | CCGGTAAGACCCAAACATACTCGCTTGTCCTTTAATGGTCC TTGAGAGGTGAAGAATACGACCACCGGGTGGTAAAATGG |
| TNNAND92_rev | CTAGCCATTTTACCACCCGGTGGTCGTATTCTTCACCTCTCA AGGACCATTAAAGGACAAGCGAGTATGTTTGGGTCTTA |
| TNNAND93_fw | CCGGTTAAATGCAAACATACTCGCTTGTCCTTTAATGGTCCT TGAGAGGTGAAGAATACGACCACCGCTGCGGAAAATGG |
| TNNAND93_rev | CTAGCCATTTTCCGCAGCGGTGGTCGTATTCTTCACCTCTC AAGGACCATTAAAGGACAAGCGAGTATGTTTGCATTTAA |

### Table S2 - TNNAND1/8 Pool Sequences

| ssDNA Oligonucleotide | Sequence (5' → 3') |
|---|---|
| TNNAND1_N6 | TCCAAGCTAGATCTACCGGTGCCTAAAACATANNNGCTT GTCCTTTAATGGTCCNNNAGAGGTGAAGAATACGACCAC CTAGGCAAAATGGCTAGCAAAGGAGA |
| TNNAND8_T1(N10) | TCCAAGCTAGATCTACCGGTNNNNNAAACATACTCGCTT GTCCTTTAATGGTCCTTGAGAGGTGAAGAATACGACCAC CNNNNNAAAATGGCTAGCAAAGGAGA |
| TNNAND8_T2(N10) | TCCAAGCTAGATCTACCGGTNNNNNCAAACATACTCGCT TGTCCTTTAATGGTCCTTGAGAGGTGAAGAATACGACCA CCGNNNNNAAAATGGCTAGCAAAGGAGA |
| TNNAND8_T3(N10) | TCCAAGCTAGATCTACCGGTNNNNNCCAAACATACTCGC TTGTCCTTTAATGGTCCTTGAGAGGTGAAGAATACGACCA CCGGNNNNNAAAATGGCTAGCAAAGGAGA |
| TNNAND8_T4(N10) | TCCAAGCTAGATCTACCGGTNNNNNGCAAACATACTCGC TTGTCCTTTAATGGTCCTTGAGAGGTGAAGAATACGACCA CCGCNNNNNAAAATGGCTAGCAAAGGAGA |
